# Porcine reproductive and respiratory syndrome virus dissemination across pig production systems in the United States

**DOI:** 10.1101/2020.04.09.034181

**Authors:** Manuel Jara, David A. Rasmussen, Cesar A. Corzo, Gustavo Machado

## Abstract

Porcine reproductive and respiratory syndrome virus (PRRSV) remains widespread in the North American pig population. Despite improvements in virus characterization, it is unclear whether PRRSV infections are a product of viral circulation within a farm, within production systems (local) or across production systems (external). Here we examined the dissemination dynamics of PRRSV and the processes facilitating its spread within and among pig farms in three production systems. Overall, PRRSV genetic diversity declined since 2018, while phylodynamic results support frequent transmission across-production systems. We found that PRRSV dissemination occurred mostly through transmission between farms of different production companies, which were predominant for several months, especially from November until May when PRRSV tends to peak in the studied region. Within production systems, dissemination occurred mainly through regular pig flow (from sow to nursery and then to finisher farms); nevertheless, an important flux of PRRSV dissemination from finisher to sow and nursery farms highlighted the importance of downstream farms as sources of the virus. Farms at areas with pig density from 500 to 1000 pig/km^2^ and farms located at a range within 0.5 km and 0.7 km from major roads were more likely to infect by PRRSV, whereas farms at elevation between 41 and 61 meters and denser vegetation acted as dissemination barriers. Although remains a challenge, there is a need to disentangle the route of PRRSV transmission, results evidenced that dissemination among commercially unrelated pig production systems was intense, reinforcing the importance of farm proximity on PRRSV spread. Thus, consideration of farm location and their geographic characteristics may help to forecast dissemination. The understanding of PRRSV transmission routes has the potential to inform targeted strategies for its prevention and control. Further studies are needed to quantify the relative contribution of PRRSV transmission routes.

## Introduction

Porcine reproductive and respiratory syndrome virus (PRRSV) remains among the most costly diseases in North America (Neumann et al., 2005; Holtkamp et al., 2013; Pileri and Mateu, 2016). Despite the great advances in reducing the incidence of PRRSv in some regions (Rathkjen and Dall, 2017) by enhanced biosecurity (Silva et al., 2019) and immunization strategies (Corzo et al., 2010), every year approximately 20-30% of US breeding herds still become infected with PRRSV (MSHMP, 2020).

PRRS is caused by an enveloped positive single-stranded RNA virus in the family Arteriviridae, order Nidovirales (Cavanagh et al., 1990). The genome of PRRSV is ~15 kb and consists of 9 open reading frames (ORFs) that encode seven structural proteins and 14 non-structural (Dokland, 2010). The mutation rate of PRRSV hinders the effectiveness of immune response against newly introduced variant strains, especially in farms with recurrent outbreaks (Linhares et al., 2014; Sanhueza et al., 2019). Likewise, recombination between North American wild-type virus strains and modified-live vaccines represents an important mechanism in the historical evolution and spread of PRRSV (Yuan et al., 1999).

The phylodynamic patterns of PRRSV spread and dissemination among pig systems and farm types have been previously described (Shi et al., 2010; Franzo et al., 2015; Alkhamis et al., 2016; Alkhamis et al., 2017; Sun et al., 2019). While there was support for between-production system dissemination (Alkhamis et al., 2016), the relevance of transmission in the maintenance of local incidence, in comparison with within-production system circulation, remains unknown. Thus, the question of whether PRRSV circulation is the product of local dissemination between farms that are part of a common commercial network (production system) versus the result of external viral introductions from nearby infected farms of distinct production systems still remains (Arruda et al., 2018).

Bayesian phylodynamic approaches have provided a breakthrough in tracking pathogen dissemination dynamics (Baele et al., 2018; Faria et al., 2019; Hicks et al., 2020). Structured coalescent models and other phylodynamic approaches allow the estimation of relative contribution of local dissemination versus external introductions to the overall incidence of disease (Rasmussen et al., 2018; Yang et al., 2019; Hicks et al., 2020). Novel methods have been developed to integrate spatial and environmental covariates in phylodynamic analysis, to determine their influence on the spread of epidemics (Dudas et al., 2017; Fountain-Jones et al., 2018; Rasmussen et al., 2018; Dellicour et al., 2019; Yang et al., 2019). Thus far, such approaches have not been widely implemented to study the dissemination of diseases in domestic animals, except for Foot-and-Mouth virus (Duchatel et al., 2019).

In this study, we used phylodynamic methods to determine the viral dissemination rates among pig production systems to assess whether different PRRSV variants are the product of local transmission or external introductions and identified the factors that facilitate or impede PRRSV spread.

## Material and methods

### Data collection

In total, our dataset comprises 4,970 ORF-5 sequences collected from 2014 to 2019, obtained from three commercially unrelated pig production systems (here coded as A, B, and C), in the US. Production system A, B and C do not shared feed truck or traded pig among themselves. Sequences were generated through outbreak investigation and from regular surveillance activities performed by each production system and shared with the Morrison Swine Health Monitoring Program (MSHMP). Each sequence was attributed to a farm type (e.g., sow, nursery, finisher, and boar stud), geographic location, pig production system (A, B, and C), and the date when the sequence was reported.

### Preliminary phylogenetic analysis

Sequences were aligned using Mega X, available at www.megasoftware.net (Kumar et al., 2018). The recombination detection program (RDP) v5.3 was used to search for recombination within our dataset (Martin et al., 2015). The alignment was screened using five methods (BootScan, Chimaera, MaxChi, RDP, and SiScan), evidencing 63 recombinant sequences, which were removed from the downstream analysis (Table S1). To determine whether there was a sufficient temporal molecular evolutionary signal, we used TempEst v1.5 (Rambaut et al., 2016). To calculate the *P*-values associated with the phylogenetic signal analysis, we followed Murray et al. (2016) using 1,000 random permutations of the sequence sampling dates (Navascués et al., 2010). The relationship found between phylogenetic tree root-to-tip divergence and sampling dates (years) supported the molecular clock analysis for this study (*P* < 0.05).

### Discrete and continuous phylogeographical analysis

Phylogenies were estimated by Bayesian inference through Markov chain Monte Carlo (MCMC), implemented in BEAST v2.5.0 (Bouckaert et al., 2014). By using ModelFinder (Kalyaanamoorthy et al., 2017) built into IQ-Tree version 1.6.1 (Nguyen et al., 2015) Supplementary Table S2, we compared different substitution models to find the best fit for our genetic dataset. The marginal likelihood value supported the use of the general time-reversible model (GTR) with gamma-distributed rate heterogeneity plus a proportion of invariable sites (GTR+G+I) (Tavaré, 1986). To reconstruct PRRSV dissemination across systems and between farm types (i.e., sow, nursery, finisher, boar stud), we used the discrete-trait phylogeographic models implemented in BEASTv2.5.0 to reconstruct ancestral states (Bouckaert et al., 2014). To explore the most important historical dissemination routes for PRRSV spread across farms, as well as among pig systems, we used Bayesian stochastic search variable selection (BSSVS) (Lemey et al., 2009). Through BSSVS, we identified the non-zero rates of change between each pair of discrete traits (production system and farm types) based on its Bayes factor value (lower than 3). To perform this analysis, an asymmetric migration rate matrix was assumed. To infer the intensity of directional transitions (forward and backward) within the migration matrix, we used a Markov jump approach. To evaluate oversampling and possible selection bias towards location and/or farm types with disproportionate amounts of samples (i.e., sow farms), we downsampled on each system to match the number of samples between the three systems, maintaining the same proportion of farm type (system A − 89%, B =−39.5%, and C=31.4 %).

To analyze the spatiotemporal spread of PRRSV, we performed a Bayesian continuous phylogeographic analysis, where the nucleotide substitution model was similar to the discrete model mentioned above. In addition, we used an uncorrelated relaxed molecular clock model with lognormal distribution (Drummond et al., 2006). The coalescent tree prior used was Bayesian SkyGrid with covariates (Gill et al., 2016). Phylogeographic inference of ancestral locations was performed under a relaxed random clock model, considering a lognormal distribution. All analyses were run for 200 million generations, sampling every 10,000th generation and removing 10% of the chain as burn-in. To estimate the relative genetic diversity, we used skygrid which is based on a nonparametric coalescent model, and is utilized to calculate the effective population size over time (Hill and Baele, 2019). To visualize the spatiotemporal spread of PRRSV, we used Spatial Phylogenetic Reconstruction of Evolutionary Dynamics using Data-Driven Documents (D3) SPREAD3 software (Bielejec et al., 2016). To evaluate the impact generated by the imbalance distribution of sequences among systems and production types, a sensitivity analysis was performed. A subsample taken from the systems and farm types with the highest number of sequences was used to match the sample size in the systems with the least number of sequenced, thus this smaller sample was reanalyzed (Supplementary material S6).

### Spatiotemporal epidemiological statistics

To summarize PRRSV diffusion over time and space we analyzed data from each production system separately and all the sequences as a whole, using the R package “seraphim” version 1.0 (Dellicour et al., 2016). Spatiotemporal information about lineage locations was extracted from 100 input trees sampled at regular intervals from the post-burn-in posterior distribution to account for phylogenetic uncertainty (Dellicour, Rose and Pybus, 2016). Each phylogeny branch was summarized and represented as a distinct vector defined by its starting geographic location, ending geographic location and dates (Dellicour, Rose and Pybus, 2016; Dellicour et al., 2016). Each branch vector represents an independent viral lineage dissemination event (Pybus et al., 2012; Laenen et al., 2016). Each vector was assigned to a diffusion coefficient value (*D*) for subsequent visualization: as described by (Pybus et al., 2012), which accounts for the time (in years) during which each specific vector (lineage) moved from its initial location until the end location of each lineage (measured from the location of the tree root). These vectors were also used to assess the invasion velocity of the sampled lineages. To determine and compare the PRRSV diffusivity associated with the overall data and for each production system, we estimated the spatial diffusion coefficient using *D*-weighted statistics (Trovão et al., 2015) since this index is less sensitive to extreme values of short branches. These results were displayed via: i) dissemination epidemiological statistics that estimated the mean branch dissemination velocity, and ii) kernel plots showing the lineage diffusion coefficients of the mean and variation on the highest posterior density (HPD).

### Between-production systems transmission: the contribution of within-system PRRSV circulation (local) versus external introduction of virus from a commercially unrelated pig farm

We estimated the contribution of local transmission (i.e., between farms within a production system) versus external introductions (i.e., between farms across production systems) using a structured coalescent model. Due to computational demand, we used PRRSV alignment from MEGA software (Kumar et al., 2008) to produce a phylogenetic tree using PhyML version 3.0 (Guindon et al., 2010), using a substitution rate = 0.00672 (estimated), and considering an HKY Hasegawa-Kishino-Yano + Γ4 (Hasegawa et al., 1985; Yang, 1994), which was dated using Least Squares Dating (LSD) version 0.3 (To et al., 2016). To infer the contribution of local versus external introductions the migration rates between different pig production systems were estimated using the Marginal Approximation to the Structured Coalescent (MASCOT) package (Müller et al., 2017, 2018) implemented in Beast2 (Bouckaert et al., 2014). To infer the evolutionary rate from the time-calibrated tree, we used a strict clock model. Using MASCOT, we estimated the effective population size in the local and external population and the migration rates between them, which was allowed to be asymmetric. This analysis was replicated three times, each time using one of the three pig systems as local and the other two as external. We also calculated the posterior probability of each internal node in the tree being in each state (i.e., local or external) using the forwards/backwards approach implemented in MASCOT (Müller et al., 2018). All analyses were performed under 100 million MCMC runs, states were sampled every 10,000 and the first 10% of samples were discarded as burn-in using TreeAnnotator v. 2.3.0 (Rambaut and Drummond, 2016). We reconstructed the number of migration events through time using the ancestral locations reconstructed by MASCOT. A migration event was assumed to occur when the most probable location of a lineage changed between its parent and child node, with the time of the migration event assumed to be the date of the daughter node. Migration events (local and external) per month were plotted using “baltic” (https://github.com/blab/baltic) in Python Software Foundation version 3.7, available at http://www.python.org. Finally, we compared the differences in local vs external transmission between PRRSV season and off-season through two-sample Wilcoxon test (Hogg et al., 2010), PRRSV season was determined by the exponentially weighted moving average proposed by MSHMP (MSHMP, 2020). In the same way as we described in the section above, a sensitivity analysis with a subsample was evaluated for consistency of the results (Supplementary material S8).

### Factors promoting or restricting PRRSv spread

To determine the effect of different predictors on the spread of PRRSV, we analyzed the results of the continuous phylogeographic analysis using the R package “seraphim” version 1.0 (Dellicour et al., 2016), considering a continuous location approach (based on the geographic coordinate location of each pig farm) and a random walk diffusion model. Spatiotemporal information was extracted from 100 trees sampled at regular intervals from the posterior distribution, after burn-in, to account for phylogenetic uncertainty. Each phylogenetic branch was considered a vector defined by its start and end location (e.g. latitude and longitude), and its start and end dates. Statistical significance for the correlation between phylogenetic and predictive factors was tested using 100 trees generated and expressed in the form of Bayes factors (BF) (Dellicour et al., 2016). Based on the extracted spatio-temporal information contained in each phylogenetic tree described above, we calculated the distances based on predictor variables associated with every branch of each tree, which were used to determine the correlation between the duration of each branch in the phylogeny and its related predictor variable distance, including the annual temperature, annual precipitation, elevation, runoff (an index that quantity of water discharged in surface streams), soil humidity (commonly used to quantify topographic control on hydrological processes) (Sørensen et al., 2005), enhanced vegetation index (EVI), which can be used to quantify vegetation greenness, as well as the pig density, and distance to the main roads (Fig. 1 and Supplementary Table S3 for more details). Briefly, the distance between each farm (geographic coordinates) to the nearest main road was calculated based on the Euclidean distance. We calculated the statistic *Q*, which represents how much variation in lineage movement is explained with spatial heterogeneity when each predictor variable is considered (Dellicour, Rose and Pybus, 2016). Here, *Q*= R^2^_env_ - R^2^_null_ where R^2^_env_ is the coefficient of determination obtained from the regression between branch duration against predictor (factors) distances, while R^2^_null_ represents the coefficient of determination for the regression between branch duration against predictor distances, which were replaced by a null raster. Thus, the factors that were considered for further Bayes factor (BF) analysis, were the ones that obtained positive *Q* values in at least 90% of the tested trees (Jacquot et al., 2017). We approximated a BF value for all predictors tested that passed the above-mentioned *Q* statistic. Predictive factors were treated as a conductance (variables that promote the spread of the disease) or a resistance factor (variables that impede its spread). For the interpretation of BF, values between 3-20 were considered “positively” supported, values between 20-150 were considered “strong”, and values >150 were considered “ overwhelming support” (Kass and Raftery, 1995). For all variables that showed a significant support either as conductance of resistance factors (considering *Q* statistic and Bayes factor) (Supplementary Table S10), each raster was subdivided into four alternative ranges based on Jenks natural breaks, for example, for elevation raster values below 20 meters constituted the low elevation areas, from 21-40 meters medium-low elevation, 41-60 medium-high elevation and >61 high elevation (Supplementary Table S11). The same continuous phylogeographic analysis using the R package “seraphim” was applied individually for each range raster as detailed above.

**Figure 1.**
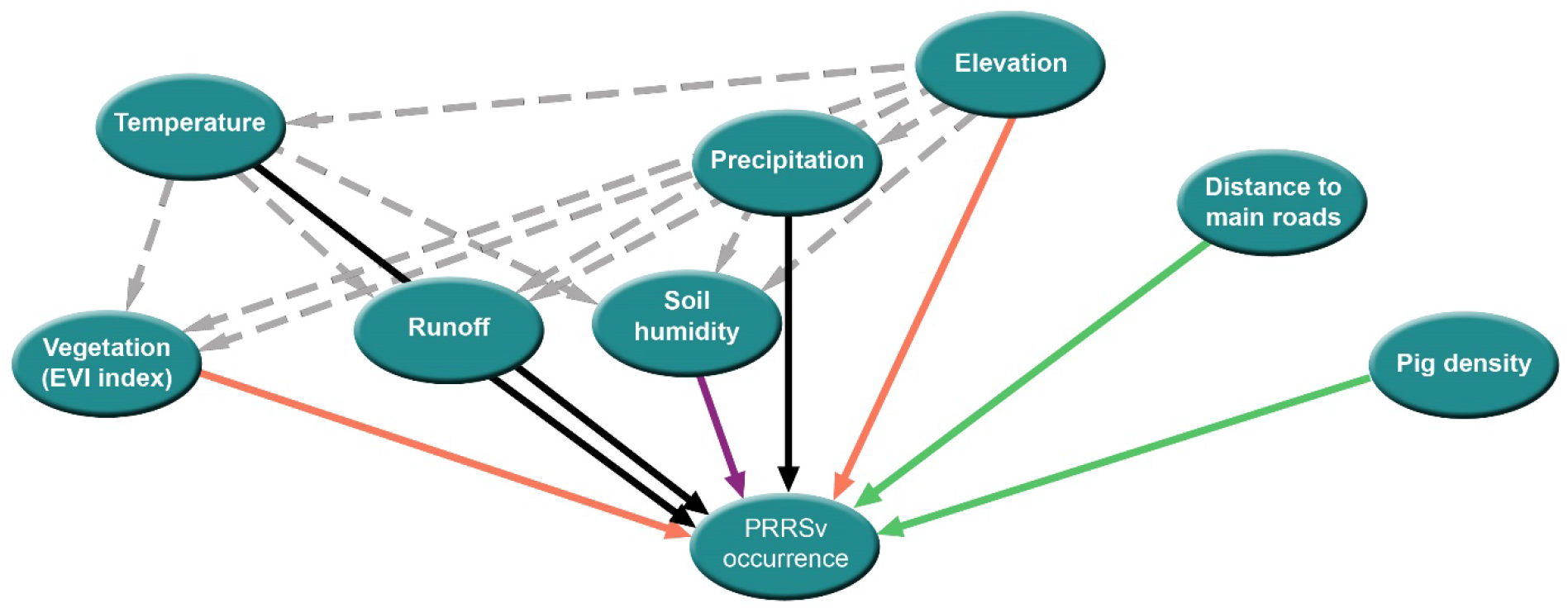
Diagram of factors promoting or restricting PRRSV spread. Arrows represent the relationship between the predictor and PRRSV occurrence based on Bayes factor analysis, where green and orange lines represent positive and negative drivers respectively. Black represent negligible support, and dashed lines show relatedness between environmental predictors affecting PRRSV through other variables.

## Results

### Study population

A total of 63 recombinant PRRSV sequences were identified and removed from the database. Most recombinants (52%) belonged to system B with the highest number of recombinants (82%) belonging to sow farms (Supplementary Table S1). Thus, 4,753 sequences remained for further discrete and continuous phylogeographic analyses. Most sequences originated from system A = 3,748 (79%), while B had 626 (13%), and C had 379 (8%). Overall, 2,958 sequences (62.2%) belonged to sow farms, 909 (19.1%) to finishers, 873 (18.4%) to nurseries, and 13 (0.3%) to boar studs (Supplementary Table S1). The nearest-neighbor distances among farms of distinct production systems had a median value of 7.6 km (1st quartile= 3.6 km and 3rd quartile= 19.3 km), the closest related production systems were A and C with median distance among all pairwise farms of 1.8 km (1st quartile= 1.2 km and 3rd quartile= 2.5 km), and finisher farms of system A and C were the closest with median of 1.9 km (1st quartile= 1.3 km, 3rd quartile= 3.2 km) (Supplementary Table S4 and S5).

### Discrete phylogeography analysis

The results of PRRSV ancestral reconstruction showed that the most likely center of origin was production system A with a root state posterior probability (RSPP) = 0.84, coming from a sow farm (RSPP = 0.75). From there, it spread heterogeneously across the other systems, unrestricted to farm types (Fig. 2A). The reconstructed phylogenetic relationship did not show evidence of any cluster related to production systems. However, we observed a clear dominance of PRRSV originating from system A over the other systems (Fig. 2A). Our sensitivity analysis on the downsample supported these results (Supplementary Table S6). The relative genetic diversity expressed as the effective population size (e.g. number of individuals contributing to a new PRRSV infection) over time showed a plateau until 2012. After that time, we captured a steep increment that lasted until approximately 2016, when relative genetic diversity decreased through June 2019 (Fig. 2B), the last month of data available for our study. The temporal pattern shown by the sampled PRRSV sequences evidenced the typical annual trend of seasonal outbreaks (Fig. 2B) (MSHMP, 2020). Phylogeographic results of system A showed that its most likely center of origin was a sow farm (RSPP = 0.47), followed by an almost equal probability for origins in finisher farm (RSPP = 0.27) and nursery farm (RSPP = 0.26), the boar stud farms were negligible (RSPP < 0.01) (Fig. 3A). In recent years, there was an evident increase in the proportion of PRRSV positive finisher farms, reaching a similar level of representation than sow farms for some clusters (Fig. 3A). SkyGrid plot revealed a relatively constant trend of relative genetic diversity until ~2011, showing a linear increase in population size, which reached its highest levels ~2016 (Fig. 3B). We observed a similar trend when we analyzed all systems together (Fig. 3B), 2015 and 2018 were the years with more samples sent to be sequenced (Fig. 3B).

**Figure 2.**
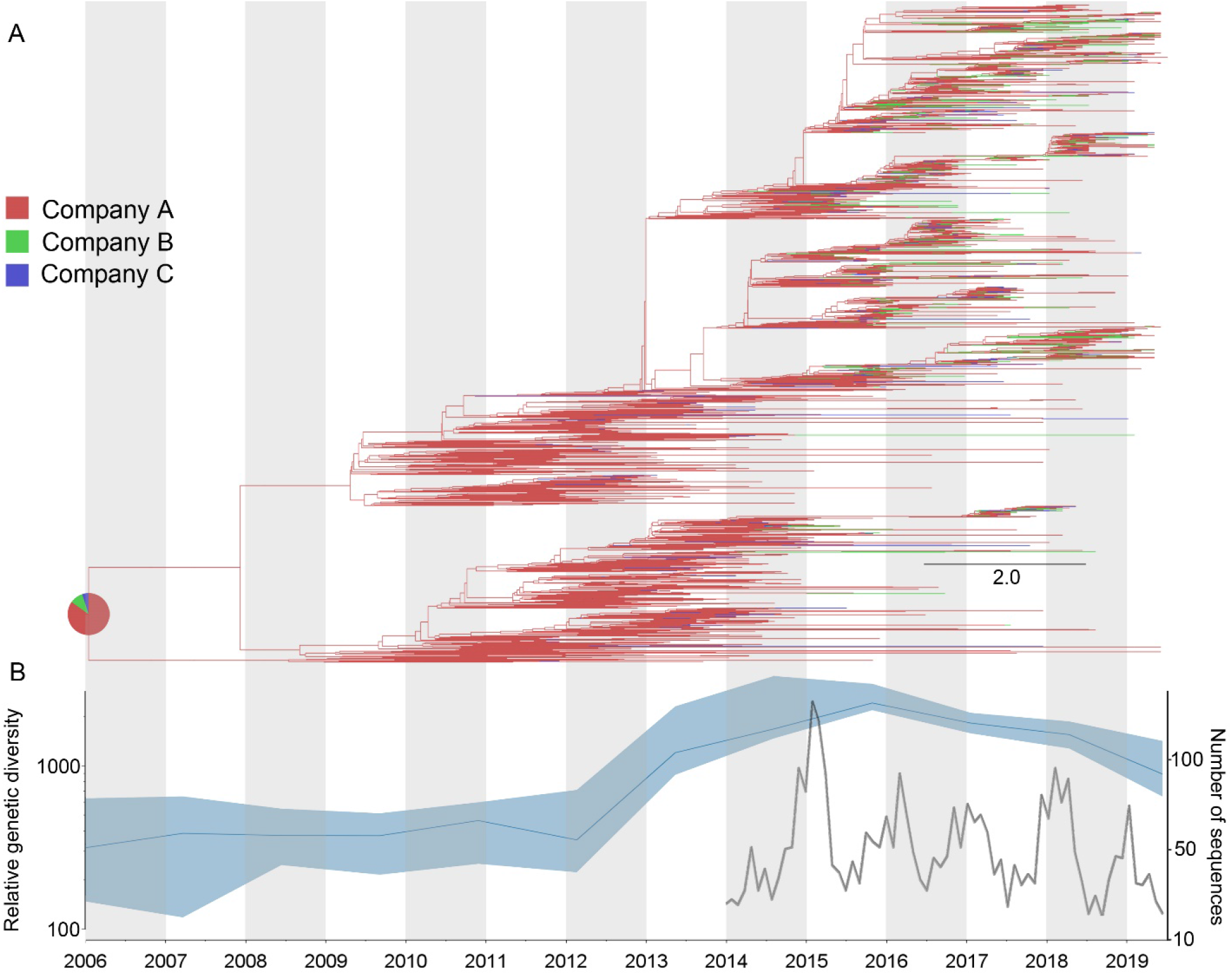
Dissemination history of PRRSV, inferred by discrete phylogeographic analysis. A) Maximum clade credibility phylogeny, colored according to pig production systems (A, B, C). The probabilities of ancestral states (inferred from the Bayesian discrete trait analysis) are shown in pie charts on each node, representing the most likely type of farms. B) Spatiotemporal patterns in the relative genetic diversity represented through the Bayesian SkyGrid plot and the number of sequences per month. The left y-axis summarizes the effective population size over time, the mean estimate is represented by the dark blue line, while the shaded light blue regions correspond to the 95% highest posterior density (HPD), and in the right y-axis shows the number of sampled sequences over time.

**Figure 3.**
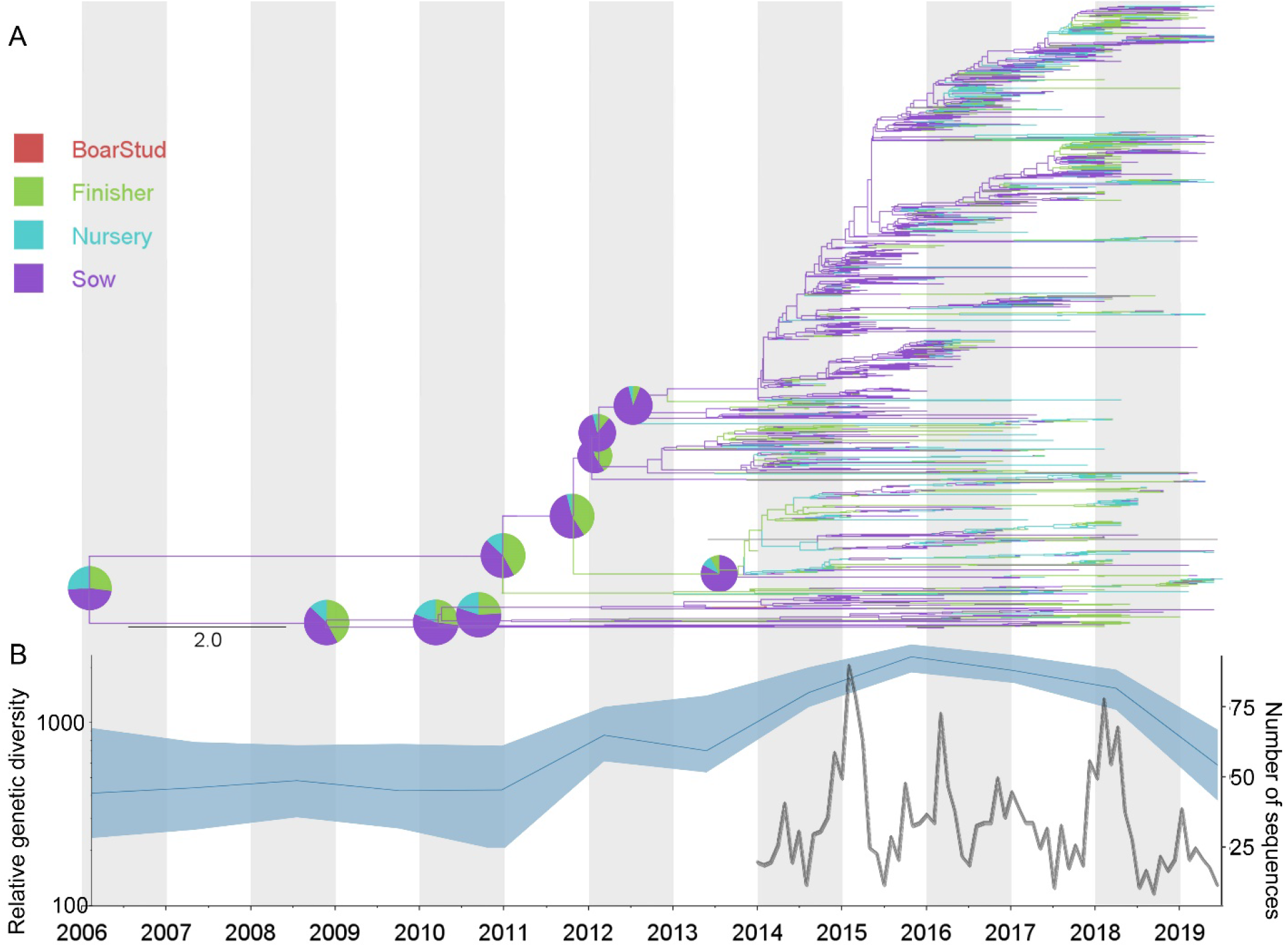
PRRSV dissemination within system A. Panel A) Maximum clade credibility phylogeny colored according to the different farm types (sow, nursery, finisher, and boar stud). Probabilities of ancestral states shown in pie charts in nodes, representing the most likely type of farm origin. Panel B) Spatiotemporal patterns in the relative genetic variation through Bayesian SkyGrid plot and number of sequences per month. The left y-axis summarizes the effective population size over time. Mean estimate is represented by the dark blue line. Shaded light blue regions correspond to the 95% highest posterior density (HPD), and in the right y-axis shows the number of sampled sequences over time.

For production system B, a sow farm (RSPP = 0.4) was also the most likely origin; however, a high probability was also linked to a nursery farm (RSPP = 0.33) (Fig. 4A). The temporal pattern shown by the effective population size was static until 2012, after which it increased rapidly until 2015-2016, when it started a linear decrease until June 2019, similar to the seasonal pattern shown by the number of sequences (Fig. 4B). As observed in systems A and B, the most likely center of origin for system C occurred inside a sow farm (RSPP = 0.52), followed by a high probability for originating in a finisher farm (RSPP = 0.4). In contrast to the other systems, we observed a low probability of origin related to a nursery farm (RSPP > 0.1) (Fig. 5A). The temporal trend in the number of sampled sequences showed the mentioned annual seasonal pattern, with peak occurrence ~2015. SkyGrid plot evidenced a more stable pattern, with higher levels of variation in effective population size during the period 2013-2014, then decreasing until 2018, when it showed a small peak and then decreased through June 2019 (Fig. 5B).

**Figure 4.**
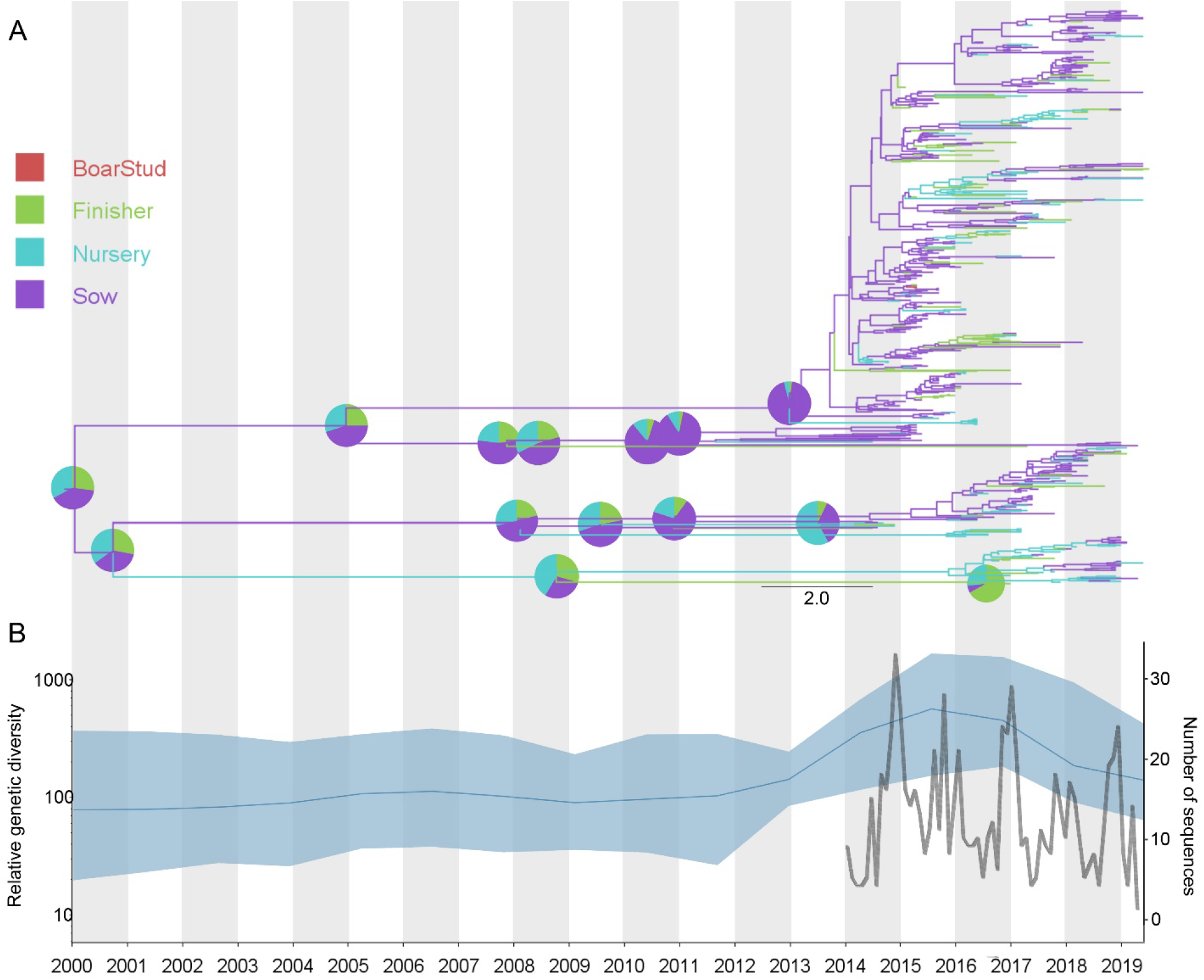
PRRSV dissemination within system B. Panel A) Maximum clade credibility phylogeny colored according to the different farm types (sow, nursery, finisher, and boar stud). Probabilities of ancestral states shown in pie charts in nodes, representing the most likely type of farm origin. Panel B) Spatiotemporal patterns in the relative genetic variation through Bayesian SkyGrid plot and number of sequences per month. The left y-axis summarizes the effective population size over time. Mean estimate is represented by the dark blue line. Shaded light blue regions correspond to the 95% highest posterior density (HPD), and in the right y-axis shows the number of sampled sequences over time.

**Figure 5.**
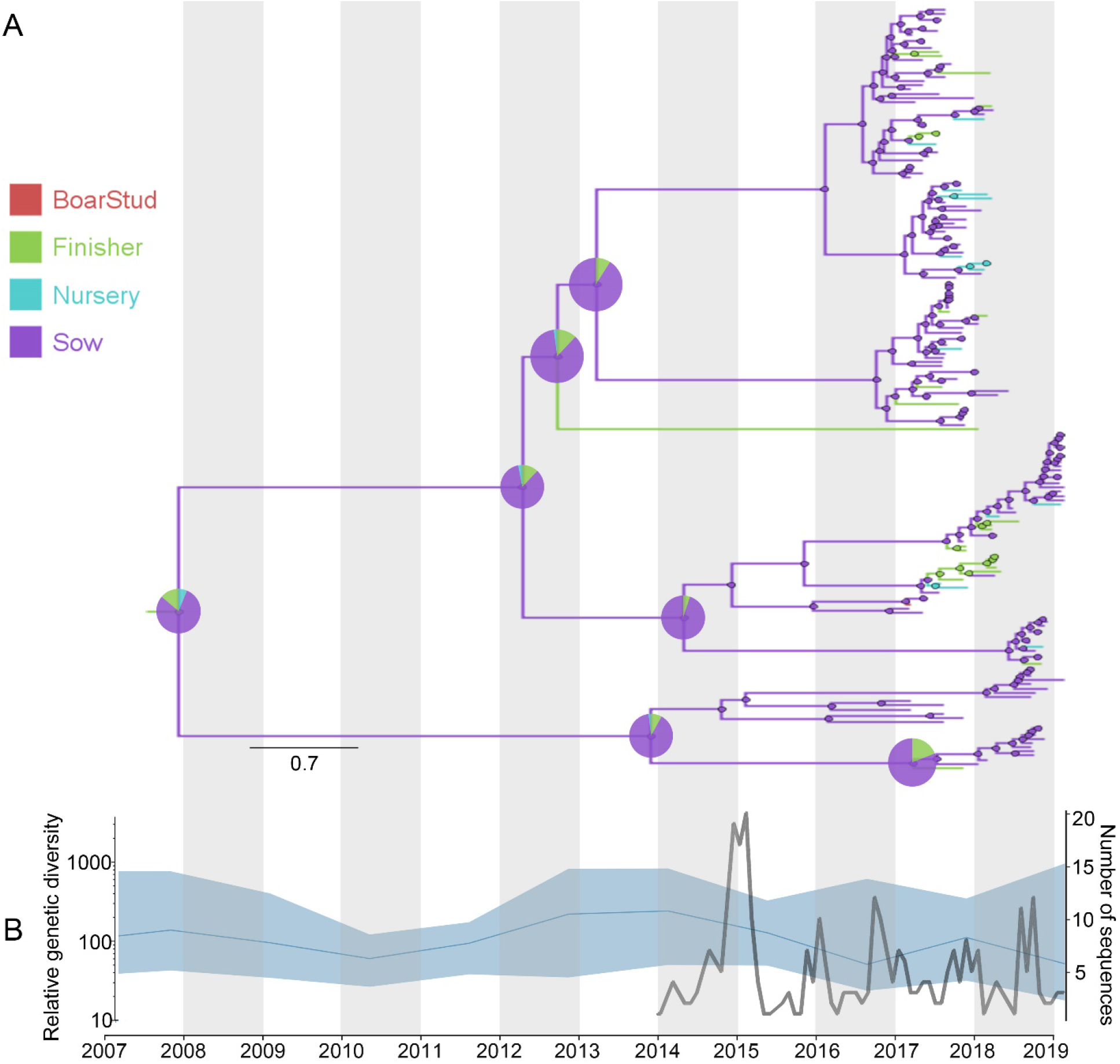
PRRSV dissemination within system C. Panel A) Maximum clade credibility phylogeny colored according to the different farm types (sow, nursery, finisher, and boar stud). Probabilities of ancestral states shown in pie charts in nodes, representing the most likely type of farm origin. Panel B) Spatiotemporal patterns in the relative genetic variation through Bayesian SkyGrid plot and number of sequences per month. The left y-axis summarizes the effective population size over time. Mean estimate is represented by the dark blue line. Shaded light blue regions correspond to the 95% highest posterior density (HPD), and in the right y-axis shows the number of sampled sequences over time.

### Phylogeographic diffusion analysis and spread statistics

The reconstructed spatiotemporal diffusion of PRRSV was characterized by continuous expansion in the geographic space, accompanied by diversification events. The comparison between each system and all samples together showed two noticeable patterns. The first one was characterized by a radial expansion over time, found when all samples were analyzed together and within samples of system A. In contrast, systems B and C evidenced a directional trend, since for both systems the spread started in the southwest region. In the case of system B, PRRSV spread to the northeast, while for system C, the spread was mainly to the north, though spread did occur in all directions (Supplementary Figs. S1-3).

The spatiotemporal analysis showed that when all sequences were analyzed together, the estimated median value for the mean branch velocity was 39 km/year (95% HPD = 31.2-61.1). This median value was lower than that shown by system A (49.2 km/year, 95% HPD= 42.9-60.4), and higher than the values observed in systems B and C (32.7 km/year, 95% HPD= 29.1-43.3), and (26.4 km/year, 95% HPD= 20.1-41.0), respectively. Estimated diffusion coefficients were asymmetric for each system and when all samples were analyzed together. System A showed the highest diffusivity (243.8 km^2^/year); however, its diffusion variation among branches was the lowest registered. The overall results (all sequences) evidenced the highest among-branch variation, while its diffusion coefficient (150 km^2^/year) was lower than system A and substantially higher than the other systems. Systems B and C showed a similar pattern in terms of diffusion coefficient variation among branches; however, B was higher than C (97 km^2^/year) and (51 km^2^/year), respectively (Supplementary Fig. S4).

### PRRSv dissemination among pig systems and farm types

Bayesian stochastic search variable selection (BSSVS) identified that the most significant disseminations were highly related to system A, evidencing strong support by a Bayes Factor score > 20. The statistical support or evidence for system A sourcing most PRRSV strains to system B, BF = 102.1 (average relative migration rate “ARMR”= 1.79), followed by dissemination from A to C, BF = 47.8 (ARMR = 0.46); of less importance was from B to A, BF= 27.4 (ARMR = 0.35). All the other disseminations between production systems were also positive, but to a lesser degree than the ones mentioned above (Table 1). In addition, a sensitivity analysis aimed to evaluate the bias related to oversampling showed that while numerically it was a difference the interpretation remains the same, but we noticed that in the subsample there was even more support for the dissemination from company B to C and company C to A (Supplementary Table S7 for more details). Our analysis reveals that when comparing PRRSV dissemination between types of farms, the most frequent routes were linked with sow farms, especially when the direction of spread was from sow to nursery (BF > 42). The highest support for this route occurred when we analyzed all systems combined (BF = 109). The second most well-supported dissemination occurred from sow to finisher farms (BF > 17.9), and as in the previous case, was best supported when we analyzed all systems together. (BF = 80.5). Likewise, the dissemination between nursery and finisher farms (in both directions) also showed positive support (BF > 6). However, all routes of dissemination that involved boar stud farms were rare events (BF < 3) (Table 2). Furthermore, the sensitivity analysis showed similar trends, where the results of the equal sample sizes these were less sharp than the one using the original dataset. However, the transmissions that involved BoarStud were all negligible (Supplementary Table S8 for more details).

**Table 1.** PRRSV dissemination between systems. Bayes factor results represent the level of support in the rates of the spread between systems, where BF > 20 is strong support, BF = 3-19 is positive support and BF< 3 is negligible.

| Origin | Destination | Average relative migration rate | Bayes factor | Interpretation |
| --- | --- | --- | --- | --- |
| A | B | $1.79 \pm 0.92$ | 102.1 | Strong support |
| A | C | $0.46 \pm 0.39$ | 47.8 | Strong support |
| B | A | $0.35 \pm 0.37$ | 27.4 | Strong support |
| B | C | $0.33 \pm 0.13$ | 7.9 | Positive support |
| C | A | $0.41 \pm 0.22$ | 14.7 | Positive support |
| C | B | $0.39 \pm 0.16$ | 10.3 | Positive support |

**Table 2.** PRRSV dissemination between types of farms. Bayes factor results represent the level of support in the rates of the spread between types of farms, where BF > 20 is strong support, BF = 3-19 is positive support and BF< 3 is negligible.

| Farm type of origin | Farm type of destination | All systems (Bayes factor) | System A (Bayes factor) | System B (Bayes factor) | System C (Bayes factor) |
| --- | --- | --- | --- | --- | --- |
| Sow | Nursery | 109 | 56.2 | 42.2 | 51.7 |
| Sow | Finisher | 80.5 | 36.6 | 28.1 | 17.9 |
| Sow | Boar stud | 0.1 | 2.2 | 0.6 | 1.1 |
| Nursery | Sow | 7.3 | 5.8 | 4.9 | 4.6 |
| Nursery | Finisher | 27.5 | 33.7 | 10.9 | 16.6 |
| Nursery | Boar stud | 1.9 | 0.1 | 0.7 | 1.3 |
| Finisher | Sow | 18.9 | 10.9 | 14.1 | 5.3 |
| Finisher | Nursery | 12 | 16.7 | 13.2 | 6.4 |
| Finisher | Boar stud | 0.2 | 1.4 | 0.1 | 0.6 |
| Boar stud | Sow | 0.5 | 1.1 | 0.3 | 0.1 |
| Boar stud | Nursery | 3.9 | 2.5 | 2.1 | 2.4 |
| Boar stud | Finisher | 0.4 | 2 | 0.8 | 1.5 |

### Within-production systems circulation (local) versus introduction of PRRSV from commercially unrelated farms (external)

We tracked the origin of the migration events leading to the current PRRSV lineages circulating among the three pig systems, to determine whether detected PRRSV strains originated from within the same system (local) or from farms of non-commercially related pig systems (external introductions). For this analysis, we considered each coalescent event in the tree as a proxy for the timing of a transmission event and then classified each event as either a local transmission event if the parent and child lineages were both reconstructed to be in the local population or an external introduction if the parent lineage resided outside the local population. Overall, results showed a clear dominance of external transmission events that were observed in all pig companies; this tendency was constant during the whole period of study (Fig. 6). System C predominantly had external introductions which peaked in 2015 with 94.7% of the transmission events, while systems A and B exhibited 82.0% in 2018 and 82.4% in 2017, respectively (Table 3). In addition, our results show that overall, the external introduction events tend to increase over time (Table 3). We also observed that coinciding with the month of higher incidence of PRRSV (MSHMP, 2020) (Fig. 6 and Supplementary Table S9). Two-sample Wilcoxon test indicated that the difference between the within farm transmission and external introductions were statistically significantly higher during PRRSV season (mean= 69.3) than the rest of the year (mean= 60.1) (Z= −9.9, *p*< 0.001).

**Figure 6.**
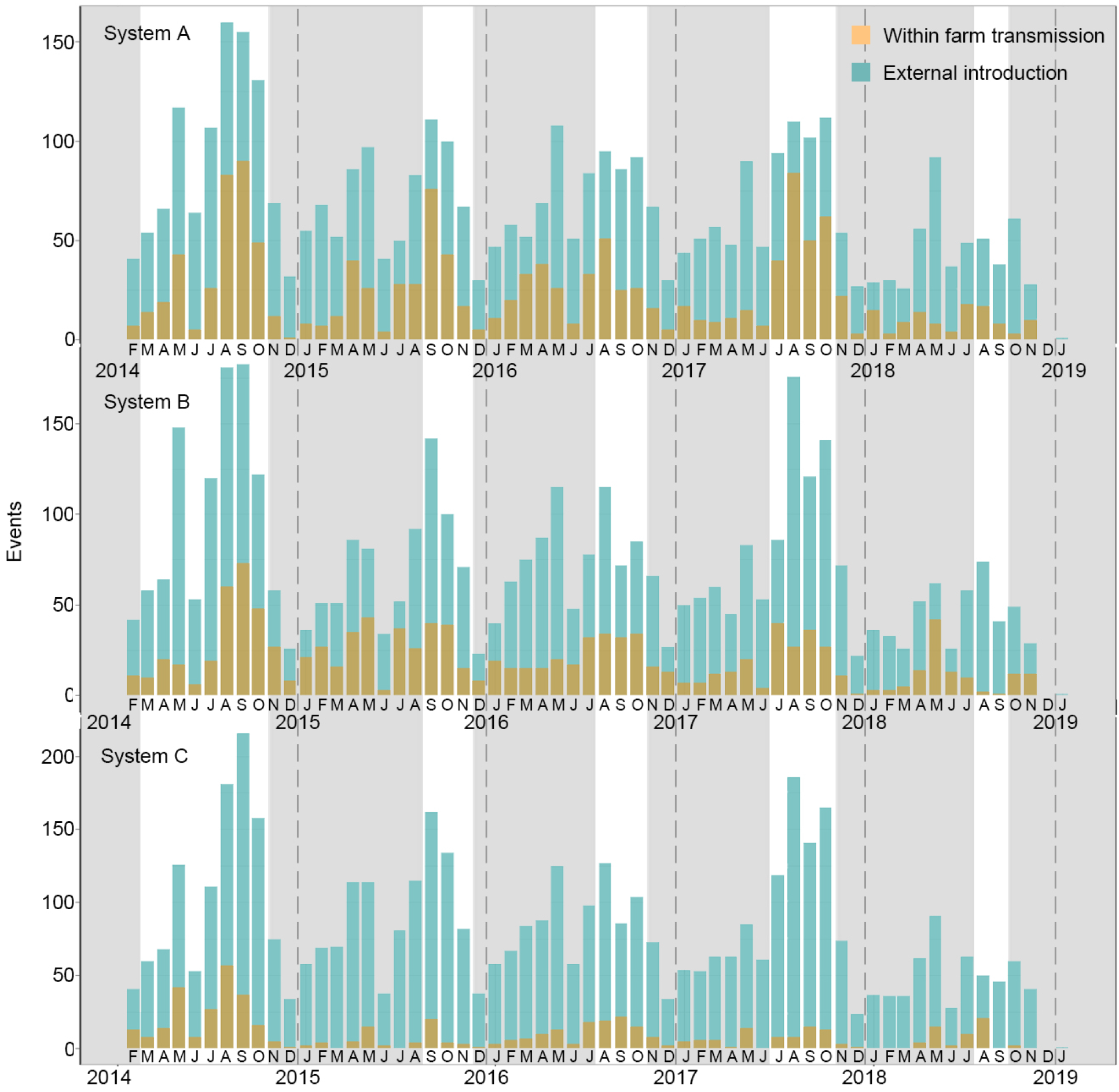
PRRSV migration events among pig systems are inferred through a structured coalescent model. Number of dissemination events originated in farms from the same pig system compared to events from farms integrated into another pig system. Grey background represents the months where the exponentially weighted moving average epidemic monitor surpassed the established PRRSv epidemic threshold based on the February 2020 MSHMP report (MSHMP, 2020), while dashed lines represent the beginning of each year.

**Table 3.** Number of transmission events that have been originated in farms that belong to the same pig system (local) compared to the events coming from farms integrated into another pig system (external).

| System | Year | Local |  | External |  | Total |
| --- | --- | --- | --- | --- | --- | --- |
|  |  | Events | % | Events | % |  |
| A | 2014 | 349 | 25.9% | 996 | 74.1% | 1345 |
|  | 2015 | 294 | 25.9% | 840 | 74.1% | 1134 |
|  | 2016 | 292 | 25.8% | 839 | 74.2% | 1131 |
|  | 2017 | 330 | 28.3% | 836 | 71.7% | 1166 |
|  | 2018 | 109 | 18.0% | 498 | 82.0% | 607 |
| B | 2014 | 299 | 22.1% | 1055 | 77.9% | 1354 |
|  | 2015 | 310 | 27.5% | 819 | 72.5% | 1129 |
|  | 2016 | 262 | 23.1% | 871 | 76.9% | 1133 |
|  | 2017 | 205 | 17.6% | 963 | 82.4% | 1168 |
|  | 2018 | 117 | 18.5% | 517 | 81.5% | 634 |
| C | 2014 | 228 | 16.9% | 1125 | 83.1% | 1353 |
|  | 2015 | 60 | 5.3% | 1075 | 94.7% | 1135 |
|  | 2016 | 126 | 11.2% | 1002 | 88.8% | 1128 |
|  | 2017 | 80 | 6.8% | 1088 | 93.2% | 1168 |
|  | 2018 | 54 | 8.9% | 551 | 91.1% | 605 |

### Factors promoting or restricting PRRSV dissemination direction

To quantify the impact of each factor on PRRSV distribution, we calculated the association between lineage duration and each covariate scaled distance through the *Q* statistic method. We found strong support (*Q* value was positive) for pig density and distance to main roads treated as conductance factors, while elevation and vegetation presented strong support as resistance factors (Supplementary Table S10 and Supplementary Figs. S5 and S6 for more details). These results also received statistical support provided by Bayes factors (BF) since we observed that farms located in areas with a high density of pig farms were more exposed to the introduction of PRRSV (BF > 12). Likewise, the distance between the farm and the nearest major road also showed a positive association with PRRSV diffusion (BF > 7). In contrast, the factors identified as negative drivers, variables that act as barriers for the spread of PRRSV, were higher elevation and denser vegetation (BF > 3) (Supplementary Table S10). With respect to the analysis of the variables with significantly supported as conductance, analyzed as ranges (Supplementary Table S11), the density of pigs ranging from 501-1.000 number of pigs/km^2^ was sufficient to facilitate the spread of PRRSV when all sequences were analyzed together and for system A, while farms at B and C were more likely infected when pig density was above 1.000 pigs/km^2^. Likewise, farms located at a distance range of 1 km to 2 km of main roads showed to favor PRRSV introductions, while when systems were analyzed in separately the dangerous distance was between 0.5 km and 1 km (Supplementary Table S11). On the other hand, for variables acting as resistance factors, our results were consistent in which at elevation above 61 meters was sufficient to slower PRRSV movement, but for system C range of 41-61 meter was sufficient. The results of vegetation index were in disagreement, while for all systems together there was no analytical support, but farms at system A, B and C an EVI index between 41 and 45 acted as a resistance range (Supplementary Table S11).

## Discussion

This analysis detailed the evolutionary epidemiology of PRRSV among commercially related and unrelated farms through continuous and discrete phylogeographic analyses. Overall, a decline in PRRSV genetic diversity was observed over time. Such findings could have different explanations, one being a rapid and efficient disseminating strain that was introduced into sow farms which consequently was transferred to growing pig sites through regular pig movement. Another potential explanation could be related to the efficacy of current interventions that local production systems have implemented, which reduced the volume and frequency of newly emerging strains, suggesting a more optimistic scenario than proposed in a similar study done in another US region (Alkhamis et al., 2017). Given the economic impact and herd health relevance of new PRRSV strain introductions, it is imperative to evaluate how often PRRSV spreads among pig production systems, determine the role of each farm type, and identify factors that may facilitate or keep PRRSV from spreading. We demonstrated that PRRSV circulation has been mostly originated by transmission from outside of each pig production system, and this tendency is more predominant during the PRRSV season (from November until May) (MSHMP, 2020).

Our findings suggest that virus transmission between pig systems is common. It is not well understood which transmission mechanism plays a role in the dissemination across systems. Airborne spread has been suggested as a potential route (Dee et al., 2009; Otake et al., 2010). However, Arruda et al., (2018, 2019) did not reach that same conclusion. Pig farm density as expected arises as a risk factor in several studies and in our study, our phylodynamic findings agree with a regional spread as an important factor for PRRSV dissemination among pig farms. PRRSV spread events appear to be relatively common in farms located in highly dense pig areas and near main roads, while farms at higher elevations and surrounded by denser vegetation were significantly less likely to report new cases of PRRSv (Arruda et al., 2017). It is important to mention that the source of our sequence data represented a limitation for our phylodynamic analysis since it was a convenience sample that can introduce bias into our analysis as the number of sequences does not directly represent all PRRSV cases circulating in this study region. There was important asymmetry in the number of sequences from boar studs, thus results of this farm type should be interpreted with caution.

### PRRSV spatiotemporal dynamics

The spatiotemporal dynamic pattern exhibited by PRRSV is the result of its transmission dynamics and how circulating strains have been affected by viral selection (Pybus and Rambaut, 2009; Volz et al., 2013). This can be observed in the shape of PRRSV phylodynamic patterns over the course of this study, which shows a clear directional selection, which can result in the loss of genetic variation (Volz et al., 2013). Based on MSHMP data (MSHMP, 2020), vaccine usage increased between 2012 and 2014, which could have impacted PRRSV genetic diversity for system A after 2012. But the increment in the number of farms using vaccines during 2014 did not show a noticeable effect on PRRSV genetic variation. However, the increment in the application of MLV’s in mid-2015 showed an undeniable positive effect on decreasing PRRSV genetic diversity (as seen in Figs. 2-5), similarly to the observations made by other authors (Sun et al., 2012; Jeong et al., 2018). However, to increase the robustness of future studies it is necessary to increase the farm coverage as well as the timeframe, not only related to the PRRSV genetic information but also to the vaccination data.

The estimated speed in which PRRSV was spreading was 39 km/year which was slightly higher than previous observations in a study that used data from one production system in another U.S. region (34 km/year) (Alkhamis et al., 2017). This difference may be related to pig farm densities, being our study region more populated with an average of 3 farms per 5 km^2^, and a more loyal representation of PRRSV dynamics since we compiled samples of more than 92% of all farms in our study area.. Consequently, there are important aspects that have to be considered while analyzing the speed at which PRRSV spreads, for example, those related with the overall pig health (i.e., absence of coinfections) and factors related with on-farm biosecurity (i.e., presence of a line of separation, cleaning, and disinfection station) (Rappole et al., 2006; Altizer et al., 2011; Lycett et al., 2019). Thus, further studies are needed to evaluate how biosecurity practices and overall pig health may contribute in slowing or accelerating PRRSV spread, results of these assessments can be used to predict the most effective biosecurity measures and optimize control strategies that go beyond PRRSV (Silva et al., 2019), not to mention current diseases that pose large-scale biosecurity threats to the U.S swine health such as African Swine Fever.

### PRRSV spread among and within farms of commercially unrelated production systems

Our analysis found relevant similarities in the spatiotemporal patterns of PRRSV spread among all farm types of all production systems (Table 1, 2), which was similar to previously reported findings that used data from one production system in the U.S. (Alkhamis et al., 2016). At farm type levels, sow farms were by far the biggest spreader (Plain and Laurence, 2003) which was expected as they wean piglets on a weekly basis and these can be transported several miles away to an off-site growing pig facility. Phylogenetic results also showed that PRRSV introductions into sow and nursery farms likely originated from finisher farms, which could be partially related to gilt development units sending replacement gilts to sow farms (Dee, 1995; Perez et al., 2015). Despite the substantial efforts to improve farms’ biosecurity and immunization through the implementation of strategies such as, sow herd air filtration, gilt acclimation, virus elimination strategies (i.e., load-close-expose and rollover) which successfully reduced the number of PRRSV outbreaks (Corzo et al., 2010; Alonso et al., 2012; Velasova et al., 2012; Alonso et al., 2013; Blanchette, 2015; Silva et al., 2019), we continue to observe significant PRRSv dissemination by indirect routes (Corzo et al., 2010; Otake et al., 2010). Therefore, it is expected that changes in immunization protocols, such as varying vaccine brands at farms of one specific system may have a serious impact on PRRSV spread.

### Within-production systems circulation (local) versus introduction of PRRSV from commercially unrelated farms (external)

Between-farm animal movement has historically been associated with disease spread (Augusta et al., 2019; Moon et al., 2019; Machado et al., 2020). Here, we used structured coalescent models to infer migration rates among commercially unrelated pig farms, calculating PRRSV dissemination not related to between-farm pig movement. Overall, PRRSV dissemination in this region has been mostly characterized by transmission between non-commercially related farms (external introductions) rather than within a production system (local). In addition, we observed that during PRRSV “season” (November-May), the external transmissions tended to be greater than during the rest of the year. While our analyses was not aimed to explore the route of transmission events or how viral lineages first entered a population (Rasmussen et al., 2018), they reinforce the importance of PRRSV collateral spread and open the question of how this dissemination is happening. Indeed, a recent work (Galvis et al., 2020) showed that for sow farms more than 50% of the between-farm transmission occurred by proximity among infected and susceptible farms, while 20% and 53% of the nurseries and finisher farms were infect by the same route, respectively. While little is known about this collateral PRRSV propagations, at some level the external introduction have been associated with contaminated vehicles that service multiple production systems (i.e., general maintenance, vaccination, cleaning or loading) (Otake et al., 2003; Dee et al., 2004), shared equipment (Pitkin et al., 2009), non-pig animal vectors (Otake et al., 2002; Schurrer et al., 2005), transport of semen (Christopher-Hennings et al., 1995) and airborne transmission (Lager and Mengeling, 2000; Trincado et al., 2004; Cho et al., 2007; Dee et al., 2009). At some degree, a contracted farm will start producing pigs under a new production system, thus it is possible that a resident virus would be introduced and become widespread within the new production system network. Knowledge gaps regarding local dissemination of PRRSV still exist, hence, studies that consider phylodynamics coupled with epidemiological models such as susceptible–infected–recovered to assess the relative contribution of between-farm PRRSV transmission routes are needed. In addition, the proposed modeling should allowed not only to answer “how it can spread” questions but also “what for”, i.e. determining what effort is required for reducing transmission (Beaunée et al., 2017).

### Factors promoting or restricting PRRSV dissemination direction

In our PRRSV dissemination models, directions of lineages revealed the potential importance of two factors that limited the spread of PRRSV. Vegetation density and high elevation, which are similar to previous observations made by Mahesh et al., (2015), Arruda et al., (2017), and (Balka et al., 2018). These authors also highlighted the important role that topography may play in mitigating the airborne spread of PRRSV. In the same way, a study on foot-and-mouth disease, which also used continuous phylogeographic analysis, identified that the presence of pure cropland or mixed land use both had a negative impact on virus diffusion (Duchatel et al., 2019). Numerous studies have shown multiple positive impacts of having vegetation barriers surrounding farms (i.e., trees, shrubs) over the airborne spread of different pathogens (Malone, 2004; Patterson et al., 2008; Van Ryswyk et al., 2019), thus it should be recognized as a possible biosecurity strategy. This study provided the first look at which amount of healthy vegetation needed to protect farms from new introductions, consistently EVI index between 41 and 45 showed to be sufficient. These values correspond to a dense tree coverage that is highly similar to the values exhibited by evergreen broadleaf forest, trees within this category are live oak (*Quercus virginiana*), laurel oak (*Quercus laurifolia*) and loblolly pine (*Pinus taeda*) (Zhang et al., 2017). Similarly, high elevated areas have proven to act as a natural topographic barrier for the local spread of pathogens (Jacquot et al., 2017), where the slope of the terrain was found to have an association with a lower PRRSV incidence (Arruda et al., 2017). We were able to identify a safe range of elevation that was consistent above 61 meter from the sea level, while this can help the local production systems, other regions of the US would need to be analyzed separately due to the differences possible elevation ranges. The identification of the main drivers associated with PRRSV dissemination could help in deciding about new farm locations, especially for systems located in highly dense pig areas (Yang et al., 2019). Farms located in areas with higher farm density were more likely to sustain new PRRSV introductions, we highlight areas in which the volume of pigs by km^2^ above 500 animals, as has been previously described by Firkins and Weigel, (2004), Arruda et al., (2017) and Alkhamis et al., (2017, 2018). In addition, our results identified distance to the main road as associated with increased chances for PRRSV circulation. This has led concern by the swine industry (Reicks, 2019), since vehicle movement has demonstrated to have a significant impact on increasing the transmission of several other diseases, for example, African Swine Fever (ASF) (Mur et al., 2012), Classical Swine Fever (CSF) (Bronsvoort et al., 2008), and for Foot and Mouth Disease (Muroga et al., 2013). Our findings indicate that farms located at 0.5 km to 0.7 km from major roads were more easily exposed to PRRSV, therefore this distance could be used to make decisions about routes used for pig movement, the enhancement of on-farm biosecurity of sites within this range of distances to major roads, and in future expansions of operations new buildings could be built beyond 0.7 km from the main road.

Here, we showed that continuous phylogeographic reconstructions represent a useful tool to describe and analyze the dissemination of PRRSV spread. Although the outcomes of phylodynamic analyses can be exploited to investigate the impact of relevant factors on lineage dissemination diffusion, it is important to note that such analyses depend on inferred viral lineage movement and thus, on the spatial distribution of the sampled sequences (Dellicour et al., 2019). Our samples represent the circulating PRRS viral population, as data comes from ~ 92% of pig farms in one highly dense pig region, therefore can be directly applied in the region. Given that most commercial pig operation in North America are similar but not the same our finding could also help the swine industry decision making on other parts of the county, conditional to data availability this analysis can be replicated at each relevant region or production system.

Our study was subject to a variety of limitations. The results of distance to main roads as a proxy for the exposure to PRRSV was inconsistent thus need to be interpreted with caution, when we analyzed all production systems together the distance of 1 km to 2 km was the only dangerous range, even though when analysis of individual systems showed that 0.7 km was enough this result need to the interpreted with caution. In the same why swine densities above > 1000 pigs/km^2^ were not supported as conductance while range of 500-1000 pigs/km^2^, in both cases distance to main road and swine density, one reason for both inconsistence results could be related to the number of farm in such range intervals. For that reason, there is an urgent need to expand the current phylodynamic framework to allow for the calculation of point estimates of the likelihood of PRRSV spread driven by one and more predictor variables more directly, which would also considered categorical variables allowing us to determine relative safe or dangerous distances to main roads, for example (Dellicour et al., 2020).

## Conclusion

Our study revealed significant asymmetries in the phylodynamic patterns among farms of three pig production systems. As a result of interventions that could be related to immunization and on-farm biosecurity, PRRSV genetic diversity has consistently declined in this region, especially in the last two years, which could explain why 2019 has had one of the lowest counts of PRRSV cases during its typical high season in the US (Sanhueza et al., 2019). Our results evidenced that PRRSV transmission across each system is most likely to originate from outside production systems. These results reiterate the importance of PRRSV farm proximity as a relevant route for virus dissemination. We highlight characteristics of locations to which PRRSV is more easily dispersed: highly dense pig population and proximity to main roads. On the protective side, higher elevation and more vegetation would be preferred. Thus, reinforcement of biosecurity may be the best tool. However, it raises the question of how much and which biosecurity practices have relevant impact on PRRSV spread. Answering this question requires further studies capable of quantifying how biosecurity practices impede or reduce PRRSV dissemination.

## Acknowledgements

We acknowledge the Department of Population Health and Pathobiology: North Carolina State University provided startup funds for Dr. Machado and CVM for the intramural grant which supported Dr. Jara. This work was also supported by Critical Agricultural Research and Extension 2019-68008-29910 from the USDA National Institute of Food and Agriculture. The Morrison Swine Health Monitoring Project is a Swine Health Information Center funded project. Authors would like to acknowledge participating systems and veterinarians.

## Authors’ contributions

MJ and GM conceived the paper ideas. MJ, DR, and GM participated in the design of the study. CC coordinated the PRRSV data collection. MJ conducted data processing and cleaning. MJ performed the phylodynamic analysis. MJ, DR, CAC, and GM wrote and edited the manuscript. All authors discussed the results and critically reviewed the manuscript.

## Ethical statement

The authors confirm the ethical policies of the journal, as noted on the journal’s author guidelines page. Since this work did not involve animal sampling nor questionnaire data collection by the researchers there was no need for ethics permits.

## Data Availability Statement

The data that support the findings of this study are not publicly available and are protected by confidential agreements, therefore, are not available.

