## supplementary for "Porcine reproductive and respiratory syndrome virus dissemination across pig production systems in the United States"

**Running title: Evolutionary epidemiology of PRRSV**

Manuel Jara^1^, David A. Rasmussen^2,3^ ,Cesar A. Corzo^4^, Gustavo Machado^1*^

^1^Department of Population Health and Pathobiology, College of Veterinary Medicine, North Carolina State University, Raleigh, NC, USA.

^2^Department of Entomology and Plant Pathology, North Carolina State University, Raleigh NC, USA

^3^Bioinformatics Research Center, North Carolina State University, Raleigh NC, USA

^4^Veterinary Population Medicine Department, College of Veterinary Medicine, University of Minnesota, 1365 Gortner Ave, St Paul, MN 55108, USA.

**Table S1.** Number of recombinant sequences by production system and farm type.

| **System** | **Farm type** | **Number of recombinants sequences** | **Number of remaining sequences** |
| --- | --- | --- | --- |
| A | Sow | 22 | 2250 |
|  | Nursery | 1 | 713 |
|  | Finisher | 2 | 776 |
|  | Boar stud | 0 | 9 |
| B | Sow | 26 | 409 |
|  | Nursery | 6 | 130 |
|  | Finisher | 1 | 85 |
|  | Boar stud | 0 | 2 |
| C | Sow | 3 | 299 |
|  | Nursery | 2 | 30 |
|  | Finisher | 0 | 48 |
|  | Boar stud | 0 | 2 |

**Table S2.** Model selection results based on best-fit molecular clock model. The model that appears in bold represents the best model chosen according to Bayesian Information Criterion (BIC).

| **Model structure** | **df** | **AIC** | **AICc** | **BIC** |
| --- | --- | --- | --- | --- |
| JC | 1097 | 36358.5 | 2445370.5 | 41187.4 |
| JC+I | 1098 | 36338.1 | 2449742.1 | 41171.4 |
| JC+G4 | 1098 | 35286.5 | 2448690.5 | 40119.8 |
| JC+I+G4 | 1099 | 35288.1 | 2453088.1 | 40125.8 |
| F81+F | 1100 | 36296.5 | 2458496.5 | 41138.6 |
| F81+F+I | 1101 | 36277.1 | 2462881.1 | 41123.6 |
| F81+F+G4 | 1101 | 35224.5 | 2461828.5 | 40071.0 |
| F81+F+I+G4 | 1102 | 35226.4 | 2466238.4 | 40077.3 |
| K2P | 1098 | 34819.1 | 2448223.1 | 39652.4 |
| K2P+I | 1099 | 34793.7 | 2452593.7 | 39631.4 |
| K2P+G4 | 1099 | 33598.1 | 2451398.1 | 38435.8 |
| K2P+I+G4 | 1100 | 33600.1 | 2455800.1 | 38442.2 |
| HKY+F | 1101 | 34758.6 | 2461362.6 | 39605.1 |
| HKY+F+I | 1102 | 34736.4 | 2465748.4 | 39587.3 |
| HKY+F+G4 | 1102 | 33553.5 | 2464565.5 | 38404.4 |
| HKY+F+I+G4 | 1103 | 33555.5 | 2468979.5 | 38410.8 |
| TNe | 1099 | 34814.6 | 2452614.6 | 39652.3 |
| TNe+I | 1100 | 34789.8 | 2456989.8 | 39631.9 |
| TNe+G4 | 1100 | 33599.1 | 2455799.1 | 38441.2 |
| TNe+I+G4 | 1101 | 33601.1 | 2460205.1 | 38447.6 |
| TN+F | 1102 | 34760.6 | 2465772.6 | 39611.5 |
| TN+F+I | 1103 | 34738.3 | 2470162.3 | 39593.6 |
| TN+F+G4 | 1103 | 33552.8 | 2468976.8 | 38408.2 |
| TN+F+I+G4 | 1104 | 33554.8 | 2473394.8 | 38414.6 |
| K3P | 1099 | 34796.6 | 2452596.6 | 39634.3 |
| K3P+I | 1100 | 34771.2 | 2456971.2 | 39613.3 |
| K3P+G4 | 1100 | 33578.7 | 2455778.7 | 38420.8 |
| K3P+I+G4 | 1101 | 33580.7 | 2460184.7 | 38427.2 |
| K3Pu+F | 1102 | 34738.6 | 2465750.6 | 39589.5 |
| K3Pu+F+I | 1103 | 34716.2 | 2470140.2 | 39571.5 |
| K3Pu+F+G4 | 1103 | 33535.8 | 2468959.8 | 38409.6 |
| K3Pu+F+I+G4 | 1104 | 33537.8 | 2473377.8 | 38397.5 |
| TPM2+F | 1102 | 34757.6 | 2465769.6 | 39608.6 |
| TPM2+F+I | 1103 | 34735.9 | 2470159.9 | 39591.2 |
| TPM2+F+G4 | 1103 | 33553.8 | 2468977.8 | 38409.1 |
| TPM2+F+I+G4 | 1104 | 33555.8 | 2473395.8 | 38415.5 |
| TPM2u+F | 1102 | 34757.6 | 2465769.6 | 39608.6 |
| TPM2u+F+I | 1103 | 34735.9 | 2470159.9 | 39591.2 |
| TPM2u+F+G4 | 1103 | 33553.8 | 2468977.8 | 38409.1 |
| TPM2u+F+I+G4 | 1104 | 33555.8 | 2473395.8 | 38415.5 |
| TPM3+F | 1102 | 34759.4 | 2465771.4 | 39610.3 |
| TPM3+F+I | 1103 | 34736.9 | 2470160.9 | 39592.2 |
| TPM3+F+G4 | 1103 | 33554.4 | 2468978.4 | 38409.8 |
| TPM3+F+I+G4 | 1104 | 33556.4 | 2473396.4 | 38416.1 |
| TPM3u+F | 1102 | 34759.4 | 2465771.4 | 39610.3 |
| TPM3u+F+I | 1103 | 34736.9 | 2470160.9 | 39592.2 |
| TPM3u+F+G4 | 1103 | 33554.4 | 2468978.4 | 38409.8 |
| TPM3u+F+I+G4 | 1104 | 33556.4 | 2473396.4 | 38416.1 |
| TIMe | 1100 | 34792.2 | 2456992.2 | 39634.3 |
| TIMe+I | 1101 | 34767.4 | 2461371.4 | 39613.9 |
| TIMe+G4 | 1101 | 33579.6 | 2460183.6 | 38426.1 |
| TIMe+I+G4 | 1102 | 33581.6 | 2464593.6 | 38432.5 |
| TIM+F | 1103 | 34740.6 | 2470164.6 | 39595.9 |
| TIM+F+I | 1104 | 34718.1 | 2474558.1 | 39577.8 |
| TIM+F+G4 | 1104 | 33535.3 | 2473375.3 | 38395.0 |
| TIM+F+I+G4 | 1105 | 33537.3 | 2477797.3 | 38401.4 |
| TIM2e | 1100 | 34816.1 | 2457016.1 | 39658.2 |
| TIM2e+I | 1101 | 34791.4 | 2461395.4 | 39637.9 |
| TIM2e+G4 | 1101 | 33601.0 | 2460205.0 | 38447.5 |
| TIM2e+I+G4 | 1102 | 33603.0 | 2464615.0 | 38453.9 |
| TIM2+F | 1103 | 34759.6 | 2470183.6 | 39614.9 |
| TIM2+F+I | 1104 | 34737.8 | 2474577.8 | 39597.5 |
| TIM2+F+G4 | 1104 | 33553.1 | 2473393.1 | 38412.8 |
| TIM2+F+I+G4 | 1105 | 33555.1 | 2477815.1 | 38419.2 |
| TIM3e | 1100 | 34812.6 | 2457012.6 | 39654.7 |
| TIM3e+I | 1101 | 34787.4 | 2461391.4 | 39633.9 |
| TIM3e+G4 | 1101 | 33594.4 | 2460198.4 | 38441.0 |
| TIM3e+I+G4 | 1102 | 33596.4 | 2464608.4 | 38447.4 |
| TIM3+F | 1103 | 34761.3 | 2470185.3 | 39616.7 |
| TIM3+F+I | 1104 | 34738.8 | 2474578.8 | 39598.6 |
| TIM3+F+G4 | 1104 | 33553.8 | 2473393.8 | 38413.5 |
| TIM3+F+I+G4 | 1105 | 33555.8 | 2477815.8 | 38419.9 |
| TVMe | 1101 | 34794.7 | 2461398.7 | 39641.2 |
| TVMe+I | 1102 | 34769.2 | 2465781.2 | 39620.1 |
| TVMe+G4 | 1102 | 33576.1 | 2464588.1 | 38427.0 |
| TVMe+I+G4 | 1103 | 33578.1 | 2469002.1 | 38433.4 |
| TVM+F | 1104 | 34735.3 | 2474575.3 | 39595.1 |
| TVM+F+I | 1105 | 34713.4 | 2478973.4 | 39577.5 |
| TVM+F+G4 | 1105 | 33535.2 | 2477795.2 | 38399.4 |
| TVM+F+I+G4 | 1106 | 33537.2 | 2482221.2 | 38405.8 |
| SYM | 1102 | 34790.3 | 2465802.3 | 39641.2 |
| SYM+I | 1103 | 34765.4 | 2470189.4 | 39620.7 |
| SYM+G4 | 1103 | 33577.0 | 2469001.0 | 38432.3 |
| SYM+I+G4 | 1104 | 33579.0 | 2473419.0 | 38438.7 |
| GTR+F | 1105 | 34737.4 | 2478997.4 | 39601.5 |
| GTR+F+I | 1106 | 34715.3 | 2483399.3 | 39583.9 |
| GTR+F+G4 | 1106 | 33534.7 | 2482218.7 | 38403.2 |
| **GTR+F+I+G4** | **1107** | **33536.7** | **2486648.7** | **38391.1** |

**Table S3.** Description and sources of the predictive factors used in this study.

| **Predictors** | **Unit** | **Source** |
| --- | --- | --- |
| Temperature | °C | <https://neo.sci.gsfc.nasa.gov/> |
| Precipitation | mm | <https://climate.northwestknowledge.net/TERRACLIMATE/index_directDownloads.php> |
| Elevation | m | <https://www.usgs.gov/> |
| Vegetation (EVI) | EVI index | <https://modis.gsfc.nasa.gov/> |
| Soil humidity | mm | [http://worldgrids.org](http://worldgrids.org/) |
| Runoff | mm | <https://climate.northwestknowledge.net/TERRACLIMATE/index_directDownloads.php> |
| Swine density | Ind/km^2^ | <https://dataverse.harvard.edu/dataverse/glw_3> |
| Distance to road | m | <https://catalog.data.gov/> |

**Table S4.** Descriptive statistics of the nearest neighbor distance between pig productive systems.

| From | To | Median (km) | Minimum (km) | q25 (km) | q75 (km) | Maximum (km) |
| --- | --- | --- | --- | --- | --- | --- |
| A | C | 5.5 | 0.1 | 2.8 | 26.1 | 222.6 |
| A | B | 14.4 | 0.5 | 6.0 | 33.2 | 229.4 |
| B | A | 2.7 | 0.5 | 1.6 | 5.6 | 12.4 |
| B | C | 7.4 | 0.7 | 3.6 | 24.5 | 114.9 |
| C | A | 1.8 | 0.1 | 1.2 | 2.5 | 28.8 |
| C | B | 14.1 | 0.7 | 6.3 | 23.5 | 94.1 |

**Table S5.** Descriptive statistics of the nearest neighbor distance between types of farm that belong to different pig productive systems.

| **From** | **To** | **Median (km)** | **Minimum (km)** | **q25 (km)** | **q75 (km)** | **Maximum (km)** |
| --- | --- | --- | --- | --- | --- | --- |
| A (Finisher) | A (Finisher) | 1.2 | 0.1 | 0.6 | 2.5 | 91.3 |
| A (Finisher) | A (Nursery) | 3.4 | 0.1 | 2.0 | 6.3 | 106.9 |
| A (Finisher) | A (Sow) | 5.8 | 0.1 | 3.2 | 10.2 | 106.4 |
| A (Finisher) | B (Finisher) | 15.8 | 0.5 | 6.7 | 30.4 | 216.7 |
| A (Finisher) | B (Nursery) | 16.3 | 0.6 | 8.0 | 32.2 | 236.8 |
| A (Finisher) | B (Sow) | 25.4 | 0.7 | 12.7 | 41.3 | 247.5 |
| A (Finisher) | C (Finisher) | 5.4 | 0.1 | 3.0 | 17.1 | 250.4 |
| A (Finisher) | C (Nursery) | 14.5 | 0.6 | 7.0 | 38.6 | 256.8 |
| A (Finisher) | C (Sow) | 16.5 | 0.9 | 7.7 | 30.7 | 222.2 |
| A (Nursery) | A (Finisher) | 2.1 | 0.1 | 1.4 | 4.0 | 78.6 |
| A (Nursery) | A (Nursery) | 1.9 | 0.1 | 0.5 | 4.2 | 127.6 |
| A (Nursery) | A (Sow) | 5.0 | 0.1 | 2.4 | 7.6 | 119.6 |
| A (Nursery) | B (Finisher) | 18.4 | 0.8 | 9.2 | 33.4 | 228.5 |
| A (Nursery) | B (Nursery) | 22.7 | 1.2 | 10.6 | 40.7 | 249.5 |
| A (Nursery) | B (Sow) | 33.0 | 1.2 | 15.7 | 51.3 | 260.2 |
| A (Nursery) | C (Finisher) | 5.5 | 0.6 | 2.9 | 11.7 | 219.1 |
| A (Nursery) | C (Nursery) | 12.7 | 1.0 | 5.6 | 35.2 | 216.5 |
| A (Nursery) | C (Sow) | 14.2 | 1.2 | 7.5 | 25.5 | 187.9 |
| A (Sow) | A (Finisher) | 3.3 | 0.1 | 1.6 | 7.0 | 100.6 |
| A (Sow) | A (Nursery) | 3.5 | 0.1 | 1.8 | 7.7 | 72.2 |
| A (Sow) | A (Sow) | 3.1 | 0.1 | 1.3 | 6.5 | 37.2 |
| A (Sow) | B (Finisher) | 28.9 | 0.6 | 12.2 | 82.3 | 195.5 |
| A (Sow) | B (Nursery) | 36.6 | 1.7 | 14.2 | 71.6 | 187.5 |
| A (Sow) | B (Sow) | 43.8 | 2.0 | 16.8 | 82.4 | 199.9 |
| A (Sow) | C (Finisher) | 10.9 | 0.6 | 4.9 | 43.0 | 166.9 |
| A (Sow) | C (Nursery) | 17.8 | 1.6 | 8.0 | 58.0 | 175.0 |
| A (Sow) | C (Sow) | 18.4 | 1.4 | 7.2 | 48.8 | 142.5 |
| B (Finisher) | A (Finisher) | 2.2 | 0.5 | 1.5 | 4.2 | 11.1 |
| B (Finisher) | A (Nursery) | 5.4 | 0.8 | 3.4 | 11.8 | 17.8 |
| B (Finisher) | A (Sow) | 11.7 | 0.6 | 6.3 | 17.1 | 28.9 |
| B (Finisher) | B (Finisher) | 2.1 | 0.3 | 1.1 | 3.8 | 34.7 |
| B (Finisher) | B (Nursery) | 5.7 | 0.1 | 3.3 | 7.8 | 29.5 |
| B (Finisher) | B (Sow) | 6.5 | 1.0 | 4.5 | 10.2 | 39.2 |
| B (Finisher) | C (Finisher) | 6.5 | 0.7 | 3.4 | 17.1 | 46.1 |
| B (Finisher) | C (Nursery) | 15.9 | 1.4 | 9.2 | 27.7 | 54.3 |
| B (Finisher) | C (Sow) | 20.0 | 3.1 | 11.4 | 25.7 | 52.1 |
| B (Nursery) | A (Finisher) | 2.2 | 0.6 | 1.4 | 6.2 | 13.6 |
| B (Nursery) | A (Nursery) | 7.3 | 1.2 | 3.8 | 10.4 | 16.4 |
| B (Nursery) | A (Sow) | 8.9 | 1.7 | 4.4 | 14.1 | 21.0 |
| B (Nursery) | B (Finisher) | 3.4 | 0.1 | 2.4 | 6.6 | 70.1 |
| B (Nursery) | B (Nursery) | 4.7 | 0.3 | 2.4 | 7.1 | 52.2 |
| B (Nursery) | B (Sow) | 6.3 | 1.0 | 3.4 | 9.7 | 39.7 |
| B (Nursery) | C (Finisher) | 7.1 | 0.9 | 3.4 | 27.0 | 110.0 |
| B (Nursery) | C (Nursery) | 13.7 | 1.6 | 8.1 | 36.8 | 120.8 |
| B (Nursery) | C (Sow) | 20.9 | 2.1 | 16.4 | 27.7 | 95.6 |
| B (Sow) | A (Finisher) | 6.6 | 0.7 | 2.6 | 10.0 | 33.9 |
| B (Sow) | A (Nursery) | 11.2 | 1.2 | 4.4 | 17.5 | 55.4 |
| B (Sow) | A (Sow) | 11.2 | 2.0 | 7.1 | 17.7 | 38.6 |
| B (Sow) | B (Finisher) | 5.1 | 1.0 | 2.4 | 43.5 | 81.8 |
| B (Sow) | B (Nursery) | 5.8 | 1.0 | 3.1 | 21.6 | 65.7 |
| B (Sow) | B (Sow) | 4.3 | 0.1 | 3.2 | 8.7 | 39.4 |
| B (Sow) | C (Finisher) | 26.1 | 1.9 | 5.2 | 87.4 | 127.0 |
| B (Sow) | C (Nursery) | 37.1 | 6.0 | 12.6 | 95.1 | 135.9 |
| B (Sow) | C (Sow) | 24.9 | 1.1 | 18.4 | 54.3 | 114.9 |
| C (Finisher) | A (Finisher) | 1.9 | 0.1 | 1.3 | 3.2 | 36.5 |
| C (Finisher) | A (Nursery) | 3.0 | 0.6 | 1.8 | 4.9 | 28.8 |
| C (Finisher) | A (Sow) | 6.0 | 0.6 | 3.5 | 9.8 | 37.2 |
| C (Finisher) | B (Finisher) | 14.9 | 0.7 | 7.1 | 22.8 | 94.1 |
| C (Finisher) | B (Nursery) | 14.8 | 0.9 | 5.8 | 23.4 | 113.8 |
| C (Finisher) | B (Sow) | 23.0 | 1.9 | 12.1 | 33.4 | 124.3 |
| C (Finisher) | C (Finisher) | 1.3 | 0.1 | 0.6 | 3.6 | 18.3 |
| C (Finisher) | C (Nursery) | 4.6 | 0.1 | 2.6 | 11.0 | 40.5 |
| C (Finisher) | C (Sow) | 8.4 | 0.1 | 3.7 | 15.3 | 40.5 |
| C (Nursery) | A (Finisher) | 2.5 | 0.6 | 1.5 | 3.8 | 30.2 |
| C (Nursery) | A (Nursery) | 2.7 | 1.0 | 2.1 | 4.2 | 28.8 |
| C (Nursery) | A (Sow) | 5.3 | 1.6 | 3.5 | 8.6 | 38.6 |
| C (Nursery) | B (Finisher) | 23.3 | 1.4 | 14.4 | 34.7 | 55.0 |
| C (Nursery) | B (Nursery) | 27.0 | 1.6 | 18.4 | 35.3 | 74.6 |
| C (Nursery) | B (Sow) | 36.8 | 6.0 | 28.0 | 46.4 | 85.0 |
| C (Nursery) | C (Finisher) | 2.4 | 0.1 | 1.7 | 4.6 | 9.2 |
| C (Nursery) | C (Nursery) | 4.6 | 0.1 | 1.2 | 9.0 | 27.3 |
| C (Nursery) | C (Sow) | 7.2 | 0.1 | 3.2 | 11.9 | 31.9 |
| C (Sow) | A (Finisher) | 2.5 | 0.9 | 1.9 | 3.1 | 29.9 |
| C (Sow) | A (Nursery) | 3.3 | 1.2 | 2.5 | 5.4 | 28.2 |
| C (Sow) | A (Sow) | 3.4 | 1.4 | 2.6 | 7.4 | 33.0 |
| C (Sow) | B (Finisher) | 20.1 | 3.1 | 12.9 | 44.0 | 59.7 |
| C (Sow) | B (Nursery) | 38.4 | 2.1 | 26.4 | 60.1 | 86.0 |
| C (Sow) | B (Sow) | 49.5 | 1.1 | 34.9 | 70.5 | 95.6 |
| C (Sow) | C (Finisher) | 5.0 | 0.1 | 1.5 | 7.6 | 56.7 |
| C (Sow) | C (Nursery) | 5.3 | 0.1 | 3.7 | 9.8 | 60.5 |
| C (Sow) | C (Sow) | 0.9 | 0.1 | 0.4 | 4.0 | 64.3 |

**Table S6.** PRRSV ancestral reconstruction shows the type of farms that are its most likely center of origin for each system, based on root state posterior probability (RSPP) results. Original= considering all sequences, ES= Equal number of samples per system and per type of farm (sensitivity analysis).

| **Farms types** | **All** | | **A** | | **B** | | **C** | |
| --- | --- | --- | --- | --- | --- | --- | --- | --- |
|  | **Original** | **ES** | **Original** | **ES** | **Original** | **ES** | **Original** | **ES** |
| Sow | 0.75 | 0.51 | 0.47 | 0.37 | 0.40 | 0.41 | 0.52 | 0.51 |
| Nursery | 0.13 | 0.23 | 0.26 | 0.31 | 0.33 | 0.35 | 0.09 | 0.17 |
| Finisher | 0.12 | 0.26 | 0.27 | 0.33 | 0.27 | 0.23 | 0.40 | 0.31 |
| Boar stud | < 0.01 | < 0.01 | < 0.01 | 0.00 | < 0.01 | < 0.01 | < 0.01 | < 0.01 |

**Table S7**. PRRSV dissemination between systems considering equal number of samples per pig system. Bayes factor results represent the level of support in the rates of the spread between pig systems (sensitivity analysis), where BF > 20 is strong support, BF = 3-19 is positive support and BF< 3 is negligible (sensitivity analysis).

| **System of origin** | **System of destination** | **Bayes factor** | **Interpretation** |
| --- | --- | --- | --- |
| A | B | 79.7 | Strong support |
| A | C | 41.0 | Strong support |
| B | A | 19.9 | Positive support |
| B | C | 15.5 | Positive support |
| C | A | 21.2 | Strong support |
| C | B | 18.2 | Positive support |

**Table S8.** PRRSV dissemination between types of farms, considering equal proportion of samples per each system and per each farm type (sensitivity analysis). Bayes factor results represent the level of support in the rates of the spread between types of farms, where BF > 20 is strong support, BF = 3-19 is positive support and BF< 3 is negligible (sensitivity analysis).

| **Farm type of origin** | **Farm type of destination** | **All sequences** | **System A** | **System B** | **System C** |
| --- | --- | --- | --- | --- | --- |
| Sow | Nursery | 73.1 | 54.2 | 41.3 | 38.3 |
| Sow | Finisher | 59.2 | 49.6 | 30.2 | 22.3 |
| Sow | BoarStud | 0.0 | 0.0 | 0.0 | 0.0 |
| Nursery | Sow | 10.7 | 17.7 | 11.2 | 6.1 |
| Nursery | Finisher | 9.2 | 23.1 | 16.9 | 20.0 |
| Nursery | BoarStud | 0.0 | 0.0 | 0.0 | 0.0 |
| Finisher | Sow | 21.9 | 8.1 | 7.9 | 5.2 |
| Finisher | Nursery | 19.4 | 7.3 | 6.8 | 6.5 |
| Finisher | BoarStud | 0.0 | 0.0 | 0.0 | 0.0 |
| BoarStud | Sow | 0.0 | 0.8 | 0.0 | 0.0 |
| BoarStud | Nursery | 0.0 | 0.0 | 0.0 | 0.0 |
| BoarStud | Finisher | 0.0 | 0.0 | 0.0 | 0.0 |

**Table S9.** Number of transition events that have originated in farms that belong to the same pig system (local) compared to the events coming from farms of another system (external).

| **System** | **Year** | **Month** | **Local** | **Local (%)** | **External** | **External (%)** | **Total** |
| --- | --- | --- | --- | --- | --- | --- | --- |
| A | 2014 | February | 7 | 14.6 | 41 | 85.4 | 48 |
|  | 2014 | March | 14 | 20.6 | 54 | 79.4 | 68 |
|  | 2014 | April | 19 | 22.4 | 66 | 77.6 | 85 |
|  | 2014 | May | 43 | 26.9 | 117 | 73.1 | 160 |
|  | 2014 | June | 5 | 7.2 | 64 | 92.8 | 69 |
|  | 2014 | July | 26 | 19.5 | 107 | 80.5 | 133 |
|  | 2014 | August | 83 | 34.2 | 160 | 65.8 | 243 |
|  | 2014 | September | 90 | 36.7 | 155 | 63.3 | 245 |
|  | 2014 | October | 49 | 27.2 | 131 | 72.8 | 180 |
|  | 2014 | November | 12 | 14.8 | 69 | 85.2 | 81 |
|  | 2014 | December | 1 | 3.0 | 32 | 97.0 | 33 |
|  | 2015 | January | 8 | 12.7 | 55 | 87.3 | 63 |
|  | 2015 | February | 7 | 9.3 | 68 | 90.7 | 75 |
|  | 2015 | March | 12 | 18.8 | 52 | 81.3 | 64 |
|  | 2015 | April | 40 | 31.7 | 86 | 68.3 | 126 |
|  | 2015 | May | 26 | 21.1 | 97 | 78.9 | 123 |
|  | 2015 | June | 4 | 8.9 | 41 | 91.1 | 45 |
|  | 2015 | July | 28 | 35.9 | 50 | 64.1 | 78 |
|  | 2015 | August | 28 | 25.2 | 83 | 74.8 | 111 |
|  | 2015 | September | 76 | 40.6 | 111 | 59.4 | 187 |
|  | 2015 | October | 43 | 30.1 | 100 | 69.9 | 143 |
|  | 2015 | November | 17 | 20.2 | 67 | 79.8 | 84 |
|  | 2015 | December | 5 | 14.3 | 30 | 85.7 | 35 |
|  | 2016 | January | 11 | 19.0 | 47 | 81.0 | 58 |
|  | 2016 | February | 20 | 25.6 | 58 | 74.4 | 78 |
|  | 2016 | March | 33 | 38.8 | 52 | 61.2 | 85 |
|  | 2016 | April | 38 | 35.5 | 69 | 64.5 | 107 |
|  | 2016 | May | 26 | 19.4 | 108 | 80.6 | 134 |
|  | 2016 | June | 8 | 13.6 | 51 | 86.4 | 59 |
|  | 2016 | July | 33 | 28.2 | 84 | 71.8 | 117 |
|  | 2016 | August | 51 | 34.9 | 95 | 65.1 | 146 |
|  | 2016 | September | 25 | 22.5 | 86 | 77.5 | 111 |
|  | 2016 | October | 26 | 22.0 | 92 | 78.0 | 118 |
|  | 2016 | November | 16 | 19.3 | 67 | 80.7 | 83 |
|  | 2016 | December | 5 | 14.3 | 30 | 85.7 | 35 |
|  | 2017 | January | 17 | 27.9 | 44 | 72.1 | 61 |
|  | 2017 | February | 10 | 16.4 | 51 | 83.6 | 61 |
|  | 2017 | March | 9 | 13.6 | 57 | 86.4 | 66 |
|  | 2017 | April | 11 | 18.6 | 48 | 81.4 | 59 |
|  | 2017 | May | 15 | 14.3 | 90 | 85.7 | 105 |
|  | 2017 | June | 7 | 13.0 | 47 | 87.0 | 54 |
|  | 2017 | July | 40 | 29.9 | 94 | 70.1 | 134 |
|  | 2017 | August | 84 | 43.3 | 110 | 56.7 | 194 |
|  | 2017 | September | 50 | 32.9 | 102 | 67.1 | 152 |
|  | 2017 | October | 62 | 35.6 | 112 | 64.4 | 174 |
|  | 2017 | November | 22 | 28.9 | 54 | 71.1 | 76 |
|  | 2017 | December | 3 | 10.0 | 27 | 90.0 | 30 |
|  | 2018 | January | 15 | 34.1 | 29 | 65.9 | 44 |
|  | 2018 | February | 3 | 9.1 | 30 | 90.9 | 33 |
|  | 2018 | March | 9 | 25.7 | 26 | 74.3 | 35 |
|  | 2018 | April | 14 | 20.0 | 56 | 80.0 | 70 |
|  | 2018 | May | 8 | 8.0 | 92 | 92.0 | 100 |
|  | 2018 | June | 4 | 9.8 | 37 | 90.2 | 41 |
|  | 2018 | July | 18 | 26.9 | 49 | 73.1 | 67 |
|  | 2018 | August | 17 | 25.0 | 51 | 75.0 | 68 |
|  | 2018 | September | 8 | 17.4 | 38 | 82.6 | 46 |
|  | 2018 | October | 3 | 4.7 | 61 | 95.3 | 64 |
|  | 2018 | November | 10 | 26.3 | 28 | 73.7 | 38 |
|  | 2018 | December | 0 | 0.0 | 1 | 100.0 | 1 |
| B | 2014 | February | 11 | 20.8 | 42 | 79.2 | 53 |
|  | 2014 | March | 10 | 14.7 | 58 | 85.3 | 68 |
|  | 2014 | April | 20 | 23.8 | 64 | 76.2 | 84 |
|  | 2014 | May | 17 | 10.3 | 148 | 89.7 | 165 |
|  | 2014 | June | 6 | 10.2 | 53 | 89.8 | 59 |
|  | 2014 | July | 19 | 13.7 | 120 | 86.3 | 139 |
|  | 2014 | August | 60 | 24.9 | 181 | 75.1 | 241 |
|  | 2014 | September | 73 | 28.5 | 183 | 71.5 | 256 |
|  | 2014 | October | 48 | 28.2 | 122 | 71.8 | 170 |
|  | 2014 | November | 27 | 31.8 | 58 | 68.2 | 85 |
|  | 2014 | December | 8 | 23.5 | 26 | 76.5 | 34 |
|  | 2015 | January | 21 | 36.8 | 36 | 63.2 | 57 |
|  | 2015 | February | 27 | 34.6 | 51 | 65.4 | 78 |
|  | 2015 | March | 16 | 23.9 | 51 | 76.1 | 67 |
|  | 2015 | April | 35 | 28.9 | 86 | 71.1 | 121 |
|  | 2015 | May | 43 | 34.7 | 81 | 65.3 | 124 |
|  | 2015 | June | 3 | 8.1 | 34 | 91.9 | 37 |
|  | 2015 | July | 37 | 41.6 | 52 | 58.4 | 89 |
|  | 2015 | August | 26 | 22.0 | 92 | 78.0 | 118 |
|  | 2015 | September | 40 | 22.0 | 142 | 78.0 | 182 |
|  | 2015 | October | 39 | 28.1 | 100 | 71.9 | 139 |
|  | 2015 | November | 15 | 17.4 | 71 | 82.6 | 86 |
|  | 2015 | December | 8 | 25.8 | 23 | 74.2 | 31 |
|  | 2016 | January | 19 | 32.2 | 40 | 67.8 | 59 |
|  | 2016 | February | 15 | 19.2 | 63 | 80.8 | 78 |
|  | 2016 | March | 15 | 16.7 | 75 | 83.3 | 90 |
|  | 2016 | April | 15 | 14.7 | 87 | 85.3 | 102 |
|  | 2016 | May | 20 | 14.8 | 115 | 85.2 | 135 |
|  | 2016 | June | 17 | 26.2 | 48 | 73.8 | 65 |
|  | 2016 | July | 32 | 29.1 | 78 | 70.9 | 110 |
|  | 2016 | August | 34 | 22.8 | 115 | 77.2 | 149 |
|  | 2016 | September | 32 | 30.8 | 72 | 69.2 | 104 |
|  | 2016 | October | 34 | 28.6 | 85 | 71.4 | 119 |
|  | 2016 | November | 16 | 19.5 | 66 | 80.5 | 82 |
|  | 2016 | December | 13 | 32.5 | 27 | 67.5 | 40 |
|  | 2017 | January | 7 | 12.3 | 50 | 87.7 | 57 |
|  | 2017 | February | 7 | 11.5 | 54 | 88.5 | 61 |
|  | 2017 | March | 12 | 16.7 | 60 | 83.3 | 72 |
|  | 2017 | April | 13 | 22.4 | 45 | 77.6 | 58 |
|  | 2017 | May | 20 | 19.4 | 83 | 80.6 | 103 |
|  | 2017 | June | 4 | 7.0 | 53 | 93.0 | 57 |
|  | 2017 | July | 40 | 31.7 | 86 | 68.3 | 126 |
|  | 2017 | August | 27 | 13.3 | 176 | 86.7 | 203 |
|  | 2017 | September | 36 | 22.9 | 121 | 77.1 | 157 |
|  | 2017 | October | 27 | 16.1 | 141 | 83.9 | 168 |
|  | 2017 | November | 11 | 13.3 | 72 | 86.7 | 83 |
|  | 2017 | December | 1 | 4.3 | 22 | 95.7 | 23 |
|  | 2018 | January | 3 | 7.7 | 36 | 92.3 | 39 |
|  | 2018 | February | 3 | 8.3 | 33 | 91.7 | 36 |
|  | 2018 | March | 5 | 16.1 | 26 | 83.9 | 31 |
|  | 2018 | April | 14 | 21.2 | 52 | 78.8 | 66 |
|  | 2018 | May | 42 | 31.3 | 92 | 68.7 | 134 |
|  | 2018 | June | 13 | 33.3 | 26 | 66.7 | 39 |
|  | 2018 | July | 10 | 14.7 | 58 | 85.3 | 68 |
|  | 2018 | August | 2 | 2.6 | 74 | 97.4 | 76 |
|  | 2018 | September | 1 | 2.4 | 41 | 97.6 | 42 |
|  | 2018 | October | 12 | 19.7 | 49 | 80.3 | 61 |
|  | 2018 | November | 12 | 29.3 | 29 | 70.7 | 41 |
|  | 2018 | December | 0 | 0.0 | 1 | 100.0 | 1 |
| C | 2014 | February | 13 | 23.2 | 43 | 76.8 | 56 |
|  | 2014 | March | 8 | 11.8 | 60 | 88.2 | 68 |
|  | 2014 | April | 14 | 17.1 | 68 | 82.9 | 82 |
|  | 2014 | May | 42 | 25.0 | 126 | 75.0 | 168 |
|  | 2014 | June | 8 | 13.1 | 53 | 86.9 | 61 |
|  | 2014 | July | 27 | 19.6 | 111 | 80.4 | 138 |
|  | 2014 | August | 57 | 23.9 | 181 | 76.1 | 238 |
|  | 2014 | September | 37 | 14.6 | 216 | 85.4 | 253 |
|  | 2014 | October | 16 | 9.2 | 158 | 90.8 | 174 |
|  | 2014 | November | 5 | 6.3 | 75 | 93.8 | 80 |
|  | 2014 | December | 1 | 2.9 | 34 | 97.1 | 35 |
|  | 2015 | January | 2 | 3.3 | 58 | 96.7 | 60 |
|  | 2015 | February | 4 | 5.5 | 69 | 94.5 | 73 |
|  | 2015 | March | 0 | 0.0 | 70 | 100.0 | 70 |
|  | 2015 | April | 5 | 4.2 | 114 | 95.8 | 119 |
|  | 2015 | May | 15 | 11.6 | 114 | 88.4 | 129 |
|  | 2015 | June | 2 | 5.0 | 38 | 95.0 | 40 |
|  | 2015 | July | 0 | 0.0 | 81 | 100.0 | 81 |
|  | 2015 | August | 4 | 3.4 | 115 | 96.6 | 119 |
|  | 2015 | September | 20 | 11.0 | 162 | 89.0 | 182 |
|  | 2015 | October | 4 | 2.9 | 134 | 97.1 | 138 |
|  | 2015 | November | 3 | 3.5 | 82 | 96.5 | 85 |
|  | 2015 | December | 1 | 2.6 | 38 | 97.4 | 39 |
|  | 2016 | January | 3 | 4.9 | 58 | 95.1 | 61 |
|  | 2016 | February | 6 | 8.2 | 67 | 91.8 | 73 |
|  | 2016 | March | 7 | 7.7 | 84 | 92.3 | 91 |
|  | 2016 | April | 10 | 10.2 | 88 | 89.8 | 98 |
|  | 2016 | May | 13 | 9.4 | 125 | 90.6 | 138 |
|  | 2016 | June | 3 | 4.9 | 58 | 95.1 | 61 |
|  | 2016 | July | 18 | 15.5 | 98 | 84.5 | 116 |
|  | 2016 | August | 19 | 13.0 | 127 | 87.0 | 146 |
|  | 2016 | September | 22 | 20.4 | 86 | 79.6 | 108 |
|  | 2016 | October | 15 | 12.6 | 104 | 87.4 | 119 |
|  | 2016 | November | 8 | 9.9 | 73 | 90.1 | 81 |
|  | 2016 | December | 2 | 5.6 | 34 | 94.4 | 36 |
|  | 2017 | January | 5 | 8.5 | 54 | 91.5 | 59 |
|  | 2017 | February | 6 | 10.2 | 53 | 89.8 | 59 |
|  | 2017 | March | 6 | 8.7 | 63 | 91.3 | 69 |
|  | 2017 | April | 1 | 1.6 | 63 | 98.4 | 64 |
|  | 2017 | May | 14 | 14.1 | 85 | 85.9 | 99 |
|  | 2017 | June | 0 | 0.0 | 61 | 100.0 | 61 |
|  | 2017 | July | 8 | 6.3 | 119 | 93.7 | 127 |
|  | 2017 | August | 8 | 4.1 | 186 | 95.9 | 194 |
|  | 2017 | September | 15 | 9.6 | 141 | 90.4 | 156 |
|  | 2017 | October | 13 | 7.3 | 165 | 92.7 | 178 |
|  | 2017 | November | 3 | 3.9 | 74 | 96.1 | 77 |
|  | 2017 | December | 1 | 4.0 | 24 | 96.0 | 25 |
|  | 2018 | January | 0 | 0.0 | 37 | 100.0 | 37 |
|  | 2018 | February | 0 | 0.0 | 36 | 100.0 | 36 |
|  | 2018 | March | 0 | 0.0 | 36 | 100.0 | 36 |
|  | 2018 | April | 4 | 6.1 | 62 | 93.9 | 66 |
|  | 2018 | May | 15 | 14.2 | 91 | 85.8 | 106 |
|  | 2018 | June | 2 | 6.7 | 28 | 93.3 | 30 |
|  | 2018 | July | 10 | 13.7 | 63 | 86.3 | 73 |
|  | 2018 | August | 21 | 29.6 | 50 | 70.4 | 71 |
|  | 2018 | September | 0 | 0.0 | 46 | 100.0 | 46 |
|  | 2018 | October | 2 | 3.2 | 60 | 96.8 | 62 |
|  | 2018 | November | 0 | 0.0 | 41 | 100.0 | 41 |
|  | 2018 | December | 0 | 0.0 | 1 | 100.0 | 1 |

**Table S10**. **Factors promoting or restricting predictors of PRRSV spread**. All variables were evaluated as potential conductance (C) factors if they facilitate the spread of PRRSV, or resistance (R), if they impede its dissemination. The Bayes factor (BF) supports were only reported when p (Q > 0) is at least 90% (0.9), all the variable that reached that level appears in bold. BF results represent the level of significance, where BF > 20 is strong support, BF = 3-19 is positive support, and BF < 3 is negligible.

| **System** | **Factor** | **Regression coef.** | ***Q* statistic** | **p(Q > 0)** | **BF** |
| --- | --- | --- | --- | --- | --- |
| **All** | Temperature (C) | 0.033 (0.001, 0.113) | 0.017 ( -0.081, 0.137) | 0.13 | - |
|  | Temperature (R) | 0.044 (0.015, 0.144) | 0.003 (-0.04, 0.036) | 0.68 | - |
|  | Precipitation (C) | 0.139 (0.018, 0.221) | 0.002 (-0.001, 0.004) | 0.58 | - |
|  | Precipitation (R) | 0.053 (0.012, 0.098) | 0.002 (-0.001, 0.04) | 0.67 | - |
|  | Elevation (C) | 0.02 (0.003, 0.117) | -0.091 (-0.203, 0.011) | 0.15 | - |
|  | **Elevation (R)** | **0.121 (0.07, 0.198)** | **0.01 (0.006, 0.012)** | **0.96** | **49.1** |
|  | Vegetation (C) | 0.151 (0.007, 0.188) | 0.005 (-0.016, 0.011) | 0.21 | - |
|  | **Vegetation (R)** | **0.115 (0.018, 0.189)** | **0.005 (0.001, 0.008)** | **0.97** | **4.3** |
|  | Soil humidity (C) | 0.043 (0.007, 0.136) | 0.001 (-0.024, 0.037) | 0.69 | - |
|  | Soil humidity (R) | 0.020 (0.007, 0.104) | -0.001 (-0.011, 0.000) | 0.08 | - |
|  | Runoff (C) | 0.017 (0.011, 0.113) | -0.019 (-0.104, 0.192) | 0.16 | - |
|  | Runoff (R) | 0.033 (0.001, 0.113) | 0.017 ( -0.081, 0.137) | 0.13 | - |
|  | **Pig density (C)** | **0.052 (0.022, 0.161)** | **0.011 (0.007, 0.012)** | **0.96** | **13.2** |
|  | Pig density (R) | 0.041 (0.015, 0.128) | -0.022 (-0.127, 0.203) | 0.53 | - |
|  | **Distance to roads (C)** | **0.011 (0.001, 0.183)** | **0.066 (0.014, 0.192)** | **0.94** | **11.6** |
|  | Distance to main roads (R) | 0.041 (0.012, 0.125) | -0.019 (-0.132, 0.099) | 0.09 | - |
| **A** | Temperature (C) | 0.021 (0.006, 0.099) | -0.000 (-0.002, -0.001) | 0.08 | - |
|  | Temperature (R) | 0.018 (0.005, 0.081) | -0.000 (-0.003, 0.001) | 0.12 | - |
|  | Precipitation (C) | 0.021 (0.006, 0.069) | -0.000 (-0.002, 0.001) | 0.23 | - |
|  | Precipitation (R) | 0.019 (0.005, 0.044) | 0.001 (-0.004, 0.015) | 0.58 | - |
|  | Elevation (C) | 0.019 (0.005, 0.021) | 0.000 (-0.012, 0.003) | 0.77 | - |
|  | **Elevation (R)** | **0.183 (0.148, 0.203)** | **0.023 (-0.001, 0.068)** | **0.96** | **8.4** |
|  | Vegetation (C) | 0.017 (0.006, 0.048) | 0.000 (-0.018, 0.003) | 0.68 | - |
|  | **Vegetation (R)** | **0.151 (0.015, 0.189)** | **0.001 (-0.001, 0.007)** | **0.95** | **10.1** |
|  | Soil humidity (C) | 0.146 (0.104, 0.189) | -0.006 (-0.013, 0.002) | 0.39 | - |
|  | Soil humidity (R) | 0.150 (0.111, 0.174) | -0.007 (-0.013, 0.000) | 0.25 | - |
|  | Runoff (C) | 0.018 (0.005, 0.043) | 0.001 (-0.004, 0.002) | 0.81 | - |
|  | Runoff (R) | 0.021 (0.004, 0.062) | 0.002 (-0.002, 0.003) | 0.41 | - |
|  | **Pig density (C)** | **0.098 (0.012, 0.113)** | **0.002 (0, 0.004)** | **0.97** | **12.6** |
|  | Pig density (R) | 0.018 (0.007, 0.095) | -0.002 (-0.014, 0.015) | 0.13 | - |
|  | **Distance to roads (C)** | **0.079 (0.007, 0.136)** | **0.065 (0.001, 0.139)** | **0.98** | **7.6** |
|  | Distance to main roads (R) | 0.018 (0.006, 0.075) | -0.002 (-0.019, 0.003) | 0.07 | - |
| **B** | Temperature (C) | 0.052 (0.021, 0.141) | 0.005 (-0.061, 0.052) | 0.76 | - |
|  | Temperature (R) | 0.043 (0.017, 0.119) | 0.006 (-0.086, 0.047) | 0.39 | - |
|  | Precipitation (C) | 0.048 (0.015, 0.144) | 0.003 (-0.059, 0.034) | 0.66 | - |
|  | Precipitation (R) | 0.008 (0.013, 0.121) | 0.009 (-0.075, 0.026) | 0.42 | - |
|  | Elevation (C) | 0.043 (0.017, 0.136) | 0.003 (-0.043, 0.037) | 0.28 | - |
|  | **Elevation (R)** | **0.021 (0.009, 0.112)** | **0.001 (0.001, 0.006)** | **0.97** | **15.7** |
|  | Vegetation (C) | 0.021 (0.007, 0.116) | 0.000 (-0.003, 0.004) | 0.54 | - |
|  | **Vegetation (R)** | **0.023 (0.006, 0.103)** | **0.041 (0.028, 0.046)** | **0.98** | **4.9** |
|  | Soil humidity (C) | 0.018 (0.007, 0.095) | -0.003 (-0.014, -0.001) | 0.04 | - |
|  | Soil humidity (R) | 0.021 (0.008, 0.069) | -0.002 (-0.011, -0.002) | 0.09 | - |
|  | Runoff (C) | 0.018 (0.003, 0.043) | -0.082 (-0.017, 0.055) | 0.19 | - |
|  | Runoff (R) | 0.021 (0.009, 0.126) | -0.001 (-0.012, 0.031) | 0.27 | - |
|  | **Pig density (C)** | **0.031 (0.004, 0.085)** | **0.096 (0.01, 0.127)** | **0.97** | **15.7** |
|  | Pig density (R) | 0.029 (0.007, 0.049) | 0.006 (-0.073, 0.034) | 0.55 | - |
|  | **Distance to roads (C)** | **0.098 (0.017, 0.152)** | **0.128 (-0.023, 0.192)** | **0.93** | **9.5** |
|  | Distance to main roads (R) | 0.033 (0.002, 0.116) | 0.003 (-0.001, 0.084) | 0.22 | - |
| **C** | Temperature (C) | 0.041 (0.011, 0.159) | 0.000 (-0.002, 0.001) | 0.25 | - |
|  | Temperature (R) | 0.051 (0.021, 0.169) | 0.009 (0.001, 0.011) | 0.16 | - |
|  | Precipitation (C) | 0.041 (0.014, 0.159) | 0.000 (-0.003, 0.001) | 0.56 | - |
|  | Precipitation (R) | 0.044 (0.015, 0.143) | 0.003 (-0.068, 0.034) | 0.67 | - |
|  | Elevation (C) | 0.042 (0.014, 0.105) | 0.002 (-0.042, 0.031) | 0.63 | - |
|  | **Elevation (R)** | **0.102 (0.068, 0.188)** | **0.062 (-0.008, 0.009)** | **1.00** | **4.6** |
|  | Vegetation (C) | 0.078 (0.052, 0.185) | 0.039 (-0.000, 0.045) | 0.11 | - |
|  | **Vegetation (R)** | **0.089 (0.047. 0.165)** | **0.015 (0.015, 0.019)** | **1.00** | **5.7** |
|  | Soil humidity (C) | 0.047 (0.018, 0.197) | 0.007 (-0.001, 0.032) | 0.31 | - |
|  | Soil humidity (R) | 0.041 (0.016, 0.158) | 0.003 (-0.002, 0.019) | 0.08 | - |
|  | Runoff (C) | 0.04 (0.128, 0.181) | 0.000 (-0.022, 0.043) | 0.46 | - |
|  | Runoff (R) | 0.036 (0.009, 0.164) | -0.016 (-0.039, 0.014) | 0.28 | - |
|  | **Pig density (C)** | **0.141 (0.012, 0.182)** | **0.179 (0.044, 0.224)** | **0.98** | **19.6** |
|  | Pig density (R) | 0.019 (0.003, 0.061) | -0.019 (-0.156, 0.024) | 0.08 | - |
|  | **Distance to roads (C)** | **0.041 (0.007, 0.125)** | **0.027 (0.001, 0.139)** | **0.96** | **13.6** |
|  | Distance to main roads (R) | 0.007 (0.000, 0.174) | -0.026 (-0.116, 0.099) | 0.08 | - |

**Table S11**. **Preferred range to conduct or restrict PRRSV spread.** Variable that showed significant support either as conductance of resistance factors (considering Q statistic and Bayes factor). Here the continue rasters were subdivided into range categories, from low, medium-low, medium-high and high, but for distance to road. All variables significant on it continue format were evaluated as potential conductance (C) factors if they facilitate the spread of PRRSV, or resistance (R), if they impede its dissemination. The Bayes factor (BF) supports were only reported when p (Q > 0) is at least 90% (0.9), all the variable that reached that level appears in bold. BF results represent the level of significance, where BF > 20 is strong support, BF = 3-19 is positive support, and BF < 3 is negligible.

| **System** | **Variable** | **Range** | **Regression coef.** | **Q statistic** | **p(Q>0)** | **BF** |
| --- | --- | --- | --- | --- | --- | --- |
| All | Elevation (R) | <20 | 0.012 (0.001, 0.039) | -0.022 (-0.075, -0.003) | 0.01 | 0.64 |
|  |  | 21-40 | 0.016 (0.002, 0.042) | -0.018 (-0.092, 0.016) | 0.21 | 1.22 |
|  |  | 41-60 | 0.025 (0.014, 0.069) | -0.009 (-0.085, 0.037) | 0.48 | **3.49** |
|  |  | >61 | 0.066 (0.017, 0.206) | 0.031 (0.000, 0.135) | **0.96** | **49.12** |
|  | Vegetation (R) | <35 | 0.005 (0.001, 0.016) | -0.029 (-0.107, -0.005) | 0.02 | 1.38 |
|  |  | 36-40 | 0.0153 (0.000, 0.052) | -0.019 (-0.099. 0.017) | 0.20 | 0.75 |
|  |  | 41-45 | 0.058 (0.031, 0.053) | 0.023 (-0.034, 0.105) | 0.83 | **4.18** |
|  |  | 46-52 | 0.002 (0.001, 0.009) | -0.017 (-0.098, -0.011) | 0.04 | 1.63 |
|  | Swine density (C) | <150 | 0.042 (0.014, 0.104) | 0.007 (-0.063, 0.034) | 0.75 | 2.07 |
|  |  | 151-500 | 0.075 (0.059, 0.102) | 0.044 (-0.053, 0.079) | 0.04 | 0.59 |
|  |  | 501-1000 | 0.055 (0.03, 0.082) | 0.024 (-0.029, 0.159) | **0.91** | **6.69** |
|  |  | >1001 | 0.009 (0.000, 0.021) | -0.025 (-0.11, 0.018) | 0.14 | 2.33 |
|  | Distance to main roads (C) | <500 | 0.0152 (0.004, 0.052) | -0.019 (-0.073, -0.001) | 0.03 | 1.56 |
|  |  | 501-700 | 0.034 (0.007, 0.114) | -0.007 (-0.016, 0.007) | 0.61 | **4.26** |
|  |  | 701-1000 | 0.003 (0.000, 0.023) | -0.032 (-0.112, -0.006) | 0.01 | 0.47 |
|  |  | 1001-2000 | 0.105 (0.094, 0.149) | 0.071 (0.000, 0.117) | **0.97** | **11.05** |
|  |  | >2000 | 0.057 (0.008, 0.197) | 0.022 (-0.021, 0.127) | 0.12 | 0.94 |
| A | Elevation (R) | <20 | 0.075 (0.044, 0.111) | -0.078 (-0.119, -0.039) | 0.03 | 2.35 |
|  |  | 21-40 | 0.047 (0.034, 0.064) | -0.106 (-0.145, -0.0677) | 0.03 | 1.27 |
|  |  | 41-60 | 0.107 (0.096, 0.129) | -0.047 (-0.099, 0.016) | 0.10 | 0.47 |
|  |  | >61 | 0.196 (0.175, 0.327) | 0.042 (-0.009, 0.161) | **0.94** | **4.56** |
|  | Vegetation (R) | <35 | 0.014 (0.007, 0.027) | -0.139 (-0.187, -0.093) | 0.02 | 0.54 |
|  |  | 36-40 | 0.026 (0.01, 0.046) | -0.108 (-0.159, -0.061) | 0.03 | 0.82 |
|  |  | 41-45 | 0.181 (0.133, 0.205) | 0.027 (-0.023, 0.077) | **0.93** | **5.25** |
|  |  | 46-52 | 0.153 (0.129, 0.21) | -0.000 (-0.048, 0.058) | 0.43 | **3.17** |
|  | Swine density (C) | <150 | 0.023 (0.001, 0.061) | -0.129 (-0.17, -0.061) | 0.01 | 0.03 |
|  |  | 151-500 | 0.146 (0.124, 0.164) | -0.008 (-0.057, 0.043) | 0.35 | 0.09 |
|  |  | 501-1000 | 0.235 (0.212, 0.252) | 0.083 (0.025, 0.125) | **0.98** | **24.00** |
|  |  | >1001 | 0.137 (0.12, 0.152) | -0.015 (-0.068, 0.027) | 0.12 | 0.08 |
|  | Distance to main roads (C) | <500 | 0.144 (0.113, 0.178) | -0.009 (-0.051, 0.029) | 0.27 | 0.84 |
|  |  | 501-700 | 0.052 (0.017, 0.145) | 0.018 (0.003, 0.028) | **0.98** | **6.23** |
|  |  | 701-1000 | 0.016 (0.014, 0.044) | -0.018 (-0.098, 0.007) | 0.26 | 1.56 |
|  |  | 1001-2000 | 0.126 (0.116, 0.147) | -0.028 (-0.08, 0.031) | 0.13 | 0.27 |
|  |  | >2000 | 0.166 (0.145, 0.297) | 0.012 (-0.039, 0.131) | 0.56 | 1.99 |
| B | Elevation (R) | <20 | 0.043 (0.01, 0.139) | 0.008 (-0.054, 0.095) | 0.64 | 2.12 |
|  |  | 21-40 | 0.016 (0.001, 0.07) | -0.019 (-0.081, 0.012) | 0.06 | 1.56 |
|  |  | 41-60 | 0.013 (0.001, 0.051) | -0.022 (-0.087, -0.005) | 0.00 | 0.47 |
|  |  | >61 | 0.048 (0.024, 0.115) | 0.0131 (-0.018, 0.021) | **0.91** | **5.67** |
|  | Vegetation (R) | <35 | 0.039 (0.011, 0.115) | 0.004 (-0.016, 0.018) | 0.85 | 0.61 |
|  |  | 36-40 | 0.034 (0.002, 0.141) | -0001 (-0.058, 0.094) | 0.24 | 1.73 |
|  |  | 41-45 | 0.081 (0.066, 0.116) | 0.046 (-0.037, 0.075) | **0.90** | **5.20** |
|  |  | 46-52 | 0.032 (0.007, 0.117) | -0.003 (-0.023, 0.002) | 0.77 | 0.56 |
|  | Swine density (C) | <150 | 0.042 (0.014, 0.102) | 0.007 (-0.064, 0.036) | 0.75 | 1.91 |
|  |  | 151-500 | 0.031 (0.007, 0.058) | -0.003 (-0.078, 0.036) | 0.57 | 0.96 |
|  |  | 501-1000 | 0.025 (0.015, 0.069) | -0.009 (-0.081, 0.015) | 0.43 | **4.19** |
|  |  | >1001 | 0.119 (0.11, 0.139) | 0.085 (-0.001, 0.128) | **0.96** | **9.00** |
|  | Distance to main roads (C) | <500 | 0.037 (0.015, 0.253) | -0.016 (-0.076, 0.071) | 0.38 | 1.37 |
|  |  | 501-700 | 0.072 (0.069, 0.099) | 0.038 (-0.043, 0.062) | **0.91** | **4.93** |
|  |  | 701-1000 | 0.042 (0.007, 0.135) | 0.007 (-0.007, 0.017) | 0.39 | 2.45 |
|  |  | 1001-2000 | 0.032 (0.018, 0.087) | -0.003 (-0.063, 0.031) | 0.61 | **3.97** |
|  |  | >2000 | 0.036 (0.026, 0.164) | -0.005 (-0.05, 0.002) | 0.20 | 2.10 |
| C | Elevation (R) | <20 | 0.111 (0.007, 0.151) | -0.013 (-0.057, 0.0185) | 0.10 | 2.17 |
|  |  | 21-40 | 0.017 (0.000, 0.069) | -0.038 (-0.157, 0.0169) | 0.18 | 0.37 |
|  |  | 41-60 | 0.106 (0.012, 0.274) | 0.079 (-0.034, 0.138) | **0.92** | **6.14** |
|  |  | >61 | 0.078 (0.006, 0.304) | 0.008 (-0.102, 0.031) | 0.53 | 1.70 |
|  | Vegetation (R) | <35 | 0.069 (0.009, 0.205) | 0.008 (-0.014, 0.071) | 0.16 | 0.54 |
|  |  | 36-40 | 0.031 (0.001, 0.186) | -0.03 (-0.142, 0.089) | 0.09 | 2.85 |
|  |  | 41-45 | 0.043 (0.017, 0.085) | 0.105 (-0.007, 0.141) | **0.95** | **5.67** |
|  |  | 46-52 | 0.057 (0.005, 0.219) | 0.012 (-0.027, 0.114) | 0.62 | **7.33** |
|  | Swine density (C) | <150 | 0.007 (0.000, 0.056) | -0.055 (-0.153, -0.01) | 0.02 | 0.79 |
|  |  | 151-500 | 0.007 (0.000, 0.036) | -0.054 (-0.178, 0.002) | 0.05 | 0.69 |
|  |  | 501-1000 | 0.057 (0.046, 0.088) | -0.004 (-0.126, 0.058) | 0.68 | **7.85** |
|  |  | >1001 | 0.184 (0.159, 0.234) | 0.121 (-0.014, 0.191) | **0.93** | **13.59** |
|  | Distance to main roads (C) | <500 | 0.076 (0.068, 0.0105) | 0.014 (-0.101, 0.071) | 0.75 | 2.02 |
|  |  | 501-700 | 0.073 (0.014, 0.24) | 0.042 (0.004, 0.130) | **1.00** | **7.45** |
|  |  | 701-1000 | 0.055 (0.009, 0.158) | -0.005 (-0.032, 0.012) | 0.36 | 1.27 |
|  |  | 1001-2000 | 0.033 (0.026, 0.062) | -0.029 (-0.152, 0.028) | 0.39 | 0.97 |
|  |  | >2000 | 0.007 (0.000, 0.057) | -0.054 (-0.153, 0.003) | 0.07 | 1.98 |

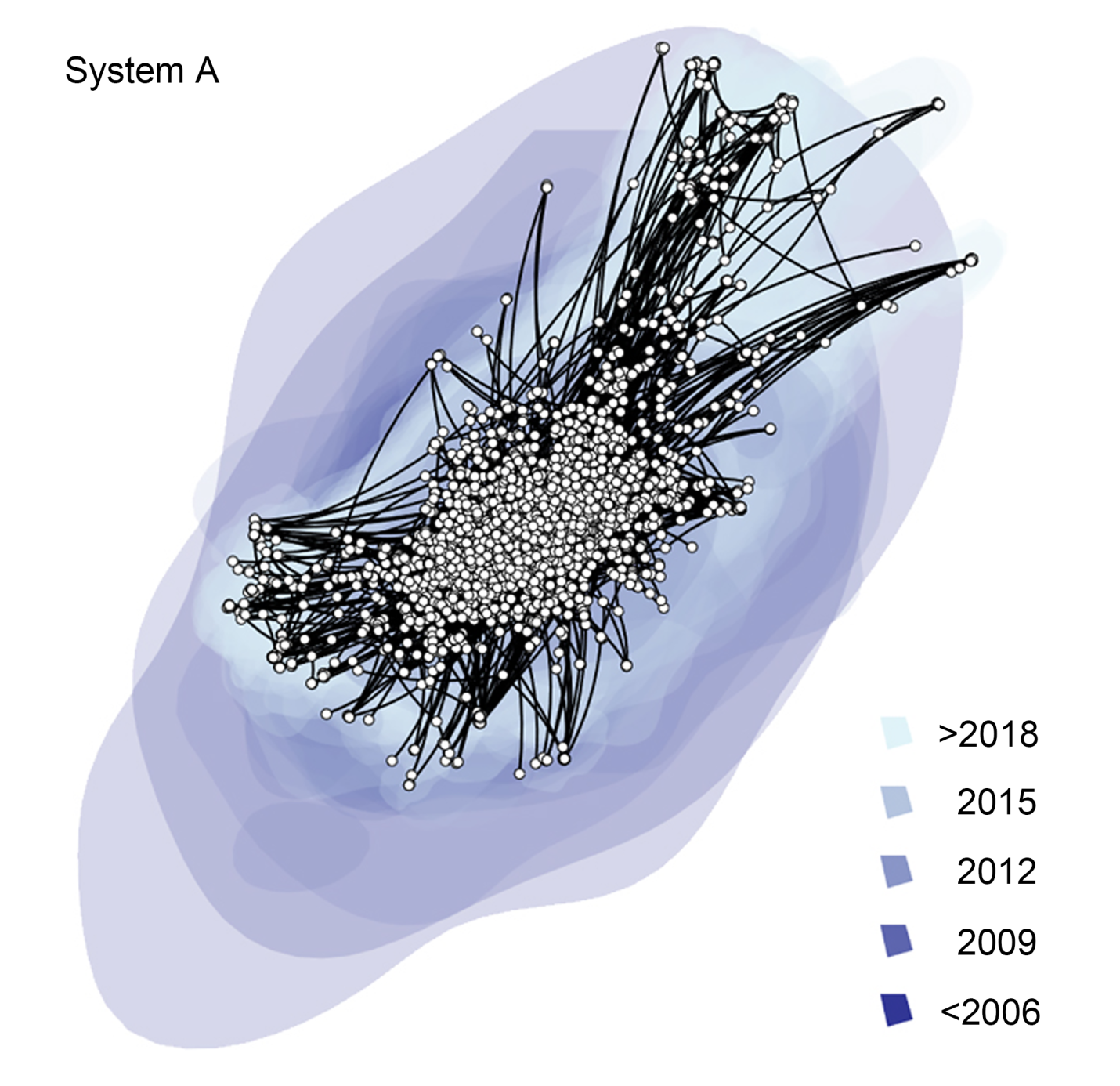

**Figure S1**. Reconstructed spatiotemporal diffusion of PRRSV on pig system A, the background color represents the age of the internal nodes, where darker blue colors represent older spread events and the white dots represent the farms and internal nodes locations.

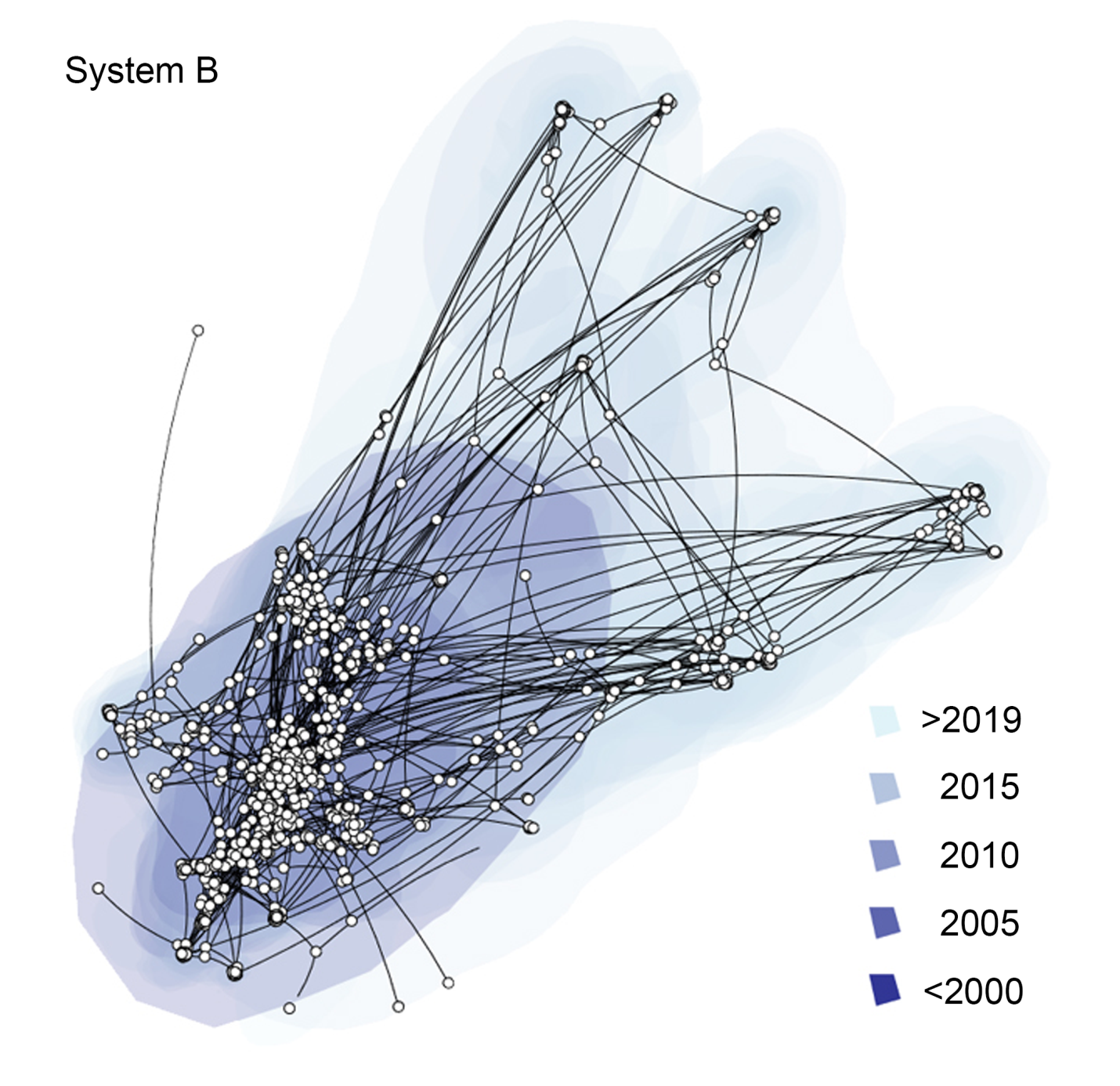

**Figure S2**. Reconstructed spatiotemporal diffusion of PRRSV on pig system B, the background color represents the age of the internal nodes, where darker blue colors represent older spread events and the white dots represent the farms and internal nodes locations.

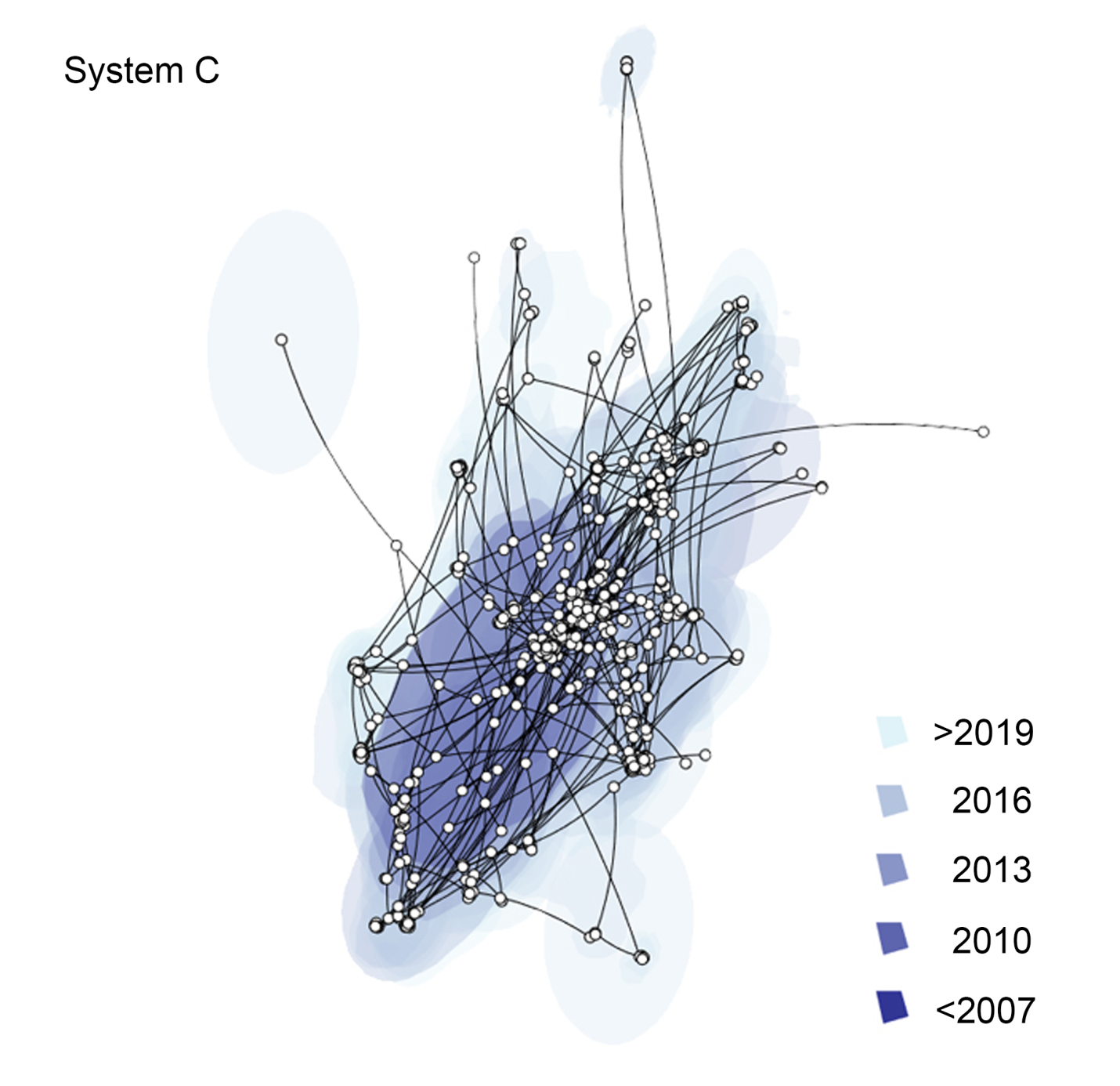

**Figure S3**. Reconstructed spatiotemporal diffusion of PRRSV on pig system C, the background color represents the age of the internal nodes, where darker blue colors represent older spread events and the white dots represent the farms and internal nodes locations.

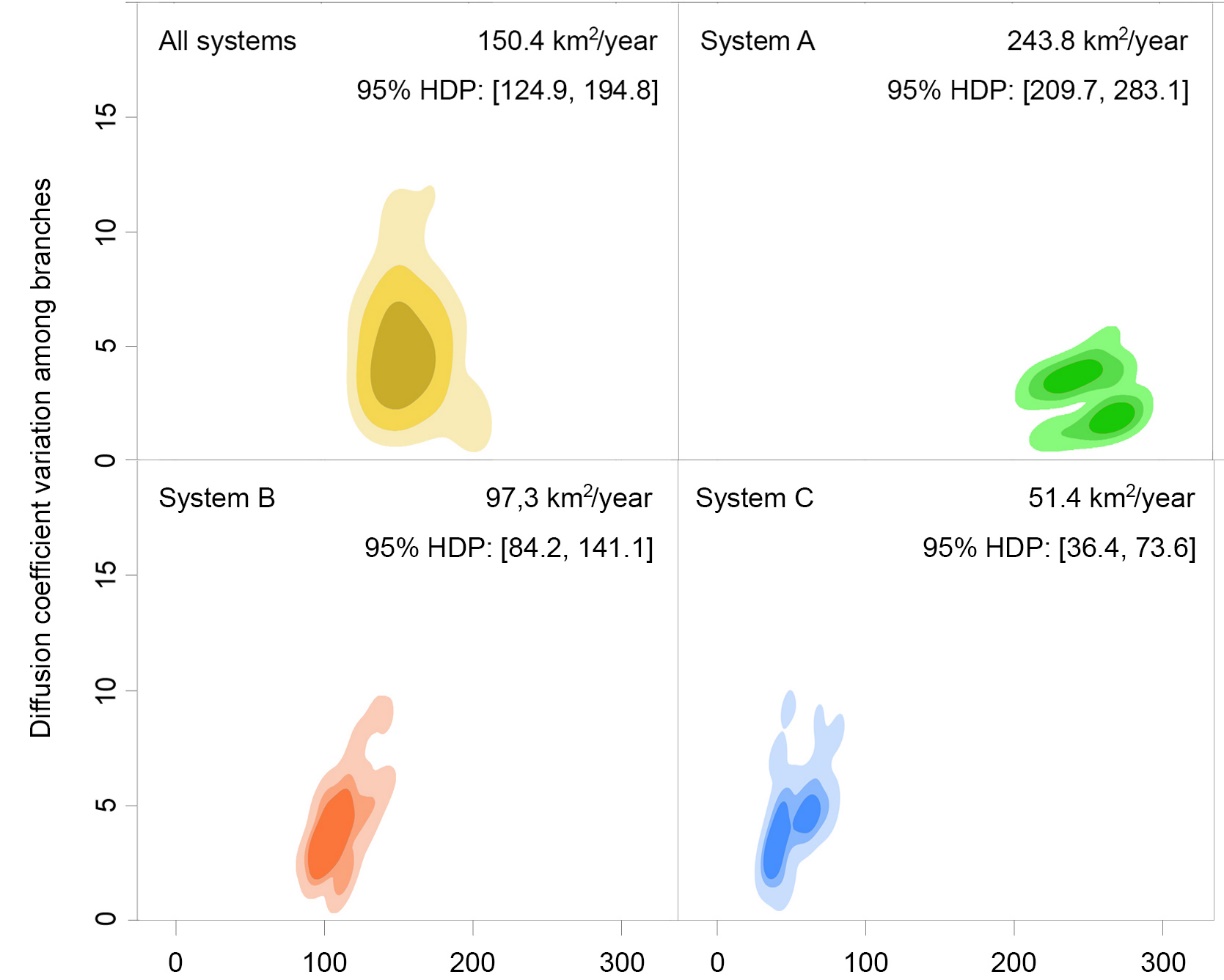

**Figure S4.** Kernel density estimates of the diffusion coefficient obtained using *the D-weighted* statistic for all PRRSV sequences and for each system. The mean diffusion coefficient among branches (x-axis) against the coefficient of variation of that value among branches (y-axis), while contours show the 50%, 75%, and 95% highest posterior density (HPD) regions via kernel density estimation. In addition, we report the median value and 95% highest posterior density intervals.

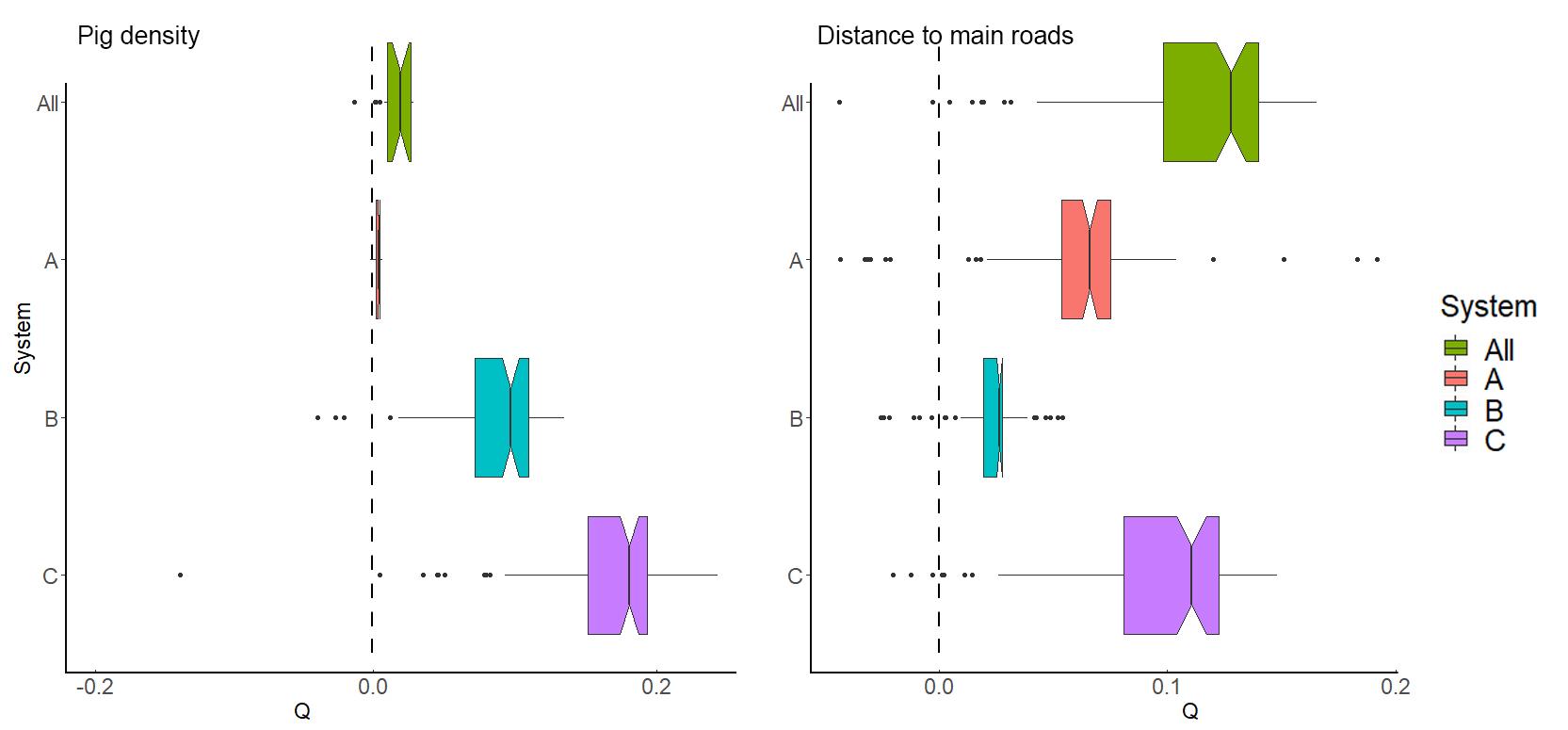

**Figure S5.** Representation of the how PRRSV variation in lineage movement is explained when spatial heterogeneity of variables of interest are treated as conductance factors through estimated Q distributions and related BF supports are reported for combinations of predictor factors for which Q > 0 in at least 90 percent of the measurements (see Table 4 for the complete results).

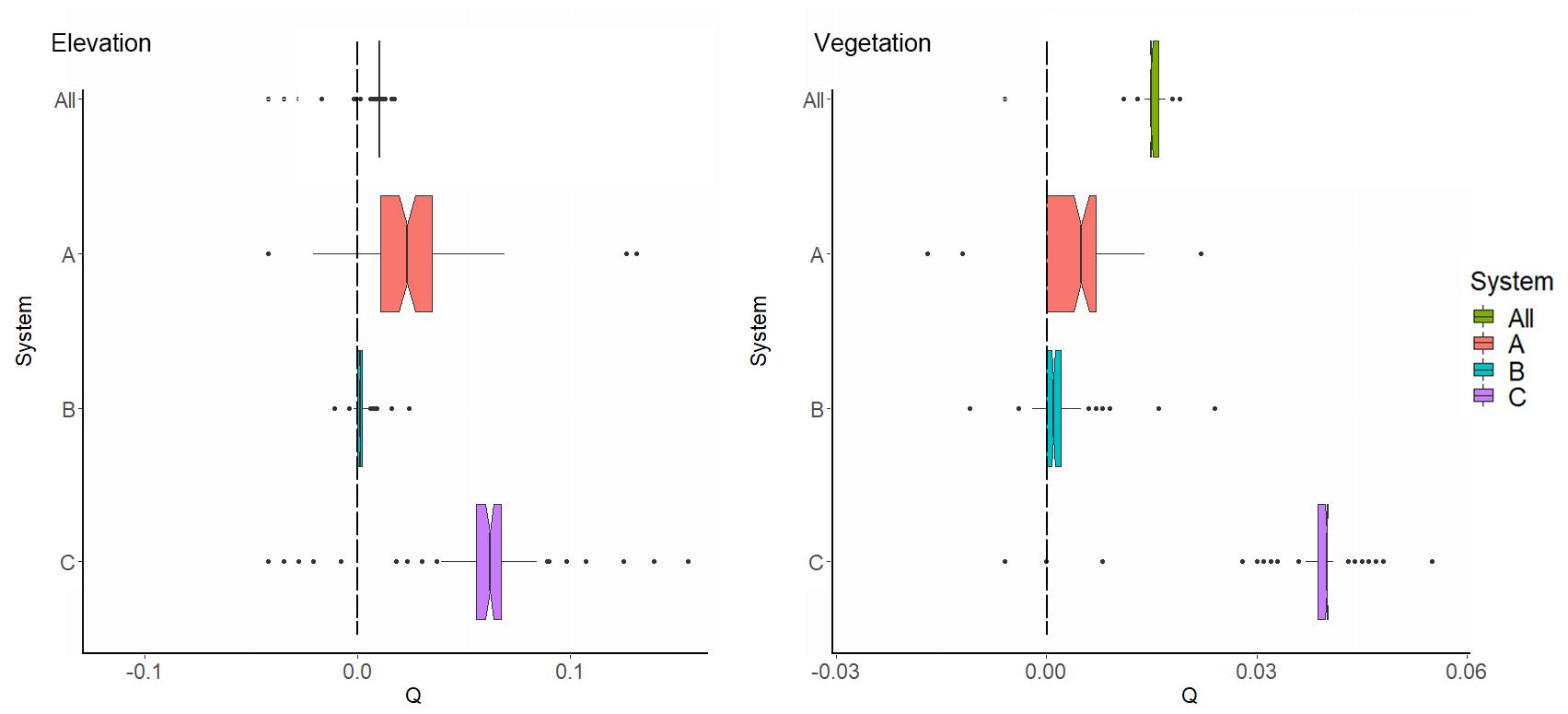

**Figure S6.** Representation of the how PRRSV variation in lineage movement is explained when spatial heterogeneity of variables of interest are treated as resistance factors through estimated Q distributions and related BF supports are reported for combinations of predictor factors for which Q > 0 in at least 90 percent of the measurements (see Table 4 for the complete results).
